# Iron limitation alters diatom carbon flow through shifts in microbiome exometabolite consumption

**DOI:** 10.64898/2026.03.16.711593

**Authors:** Naomi E. Gilbert, Nicole R. Coffey, Nathan McCall, Jeffrey A. Kimbrel, Craig A. Carlson, Elisa Halewood, Christina Ramon, Patrick Diep, Peter K. Weber, Xavier Mayali, Rene Boiteau, Rhona K. Stuart

**Affiliations:** Physical and Life Sciences Directorate, Lawrence Livermore National Laboratory, Livermore, CA, USA; Department of Chemistry, University of Minnesota, Minneapolis, Minnesota 55455, USA; Department of Ecology, Evolution, and Marine Biology, Marine Science Institute, University of California, Santa Barbara, California, USA; Bermuda Institute of Ocean Sciences, Global Futures Laboratory, Arizona State University, St George’s, Bermuda

**Author notes:** **Authors for Correspondence** (Rhona K. Stuart). Syracuse University, Department of Biology, Syracuse, NY, USA.

**Keywords:** *Phaeodactylum tricornutum*, phycosphere, nanoSIMS, nanoSIP, exometabolomics, metagenomics

## Abstract

Iron is required for photosynthesis, and thus affects biogeochemical cycling in widespread regions where its availability is limited. Turnover of aquatic photosynthetically-derived carbon is largely constrained by bacterial activity, but we lack a mechanistic understanding of how iron limitation influences this activity. We examined a bacterial enrichment community dependent on carbon from the diatom *Phaeodactylum tricornutum* to investigate how iron limitation alters the flow of carbon to bacteria, and exometabolite and community composition. Using stable isotope tracing, we quantified diatom exudate incorporation with single-cell-resolution. We identified a population of bacteria under iron limitation with high metabolic activity yet low incorporation of newly-fixed diatom carbon, indicating a shift in metabolism relative to the iron-replete control. Ultra-high-resolution exometabolomics revealed bacterial consumption of aromatics, lipid-like compounds, and purines and pyrimidines occurred under iron-limitation, when these compounds also exhibited increased exudation. We identified gene pathways for utilization of these compounds in taxa with increased abundance under iron limitation which may be responsible for carbon flow shifts. These results provide a mechanistic link between iron-limitation driven shifts in exudate composition and flow of carbon to the microbiome. This has important implications for predicting carbon flow in surface oceans and manipulating algal-bacterial interactions in engineered systems.

## Introduction

As a major trace micronutrient, iron (Fe) controls the physiology and molecular composition of microbial photoautotrophs (phytoplankton or algae), which are foundational to aquatic ecosystem processes ^1,2^. Dissolved Fe is tightly linked to the cycling of other elements such as carbon and nitrogen, and limits primary production in over one third of the surface ocean ^3–7^. The impacts of Fe deficiency on algae have been well documented and are due in part to the amount of Fe required for photosynthesis ^8^. Diatoms, which account for about 40% of marine productivity ^9^ and have biotechnological importance ^10,11^, are widely limited by Fe and dominate the phytoplankton community in regions with low Fe levels relative to other nutrients due to their diverse abilities to cope with Fe limitation ^3,12–15^. In diatoms, Fe limitation often leads to decreased growth and biomass, impaired photochemical efficiency, restructuring of photosystem II, and oxidative stress responses ^8,13,14,16^.

Heterotrophic microbes process over half of algal-derived products ^17^ and are known to be impacted by Fe limitation, which is reflected in their diverse abilities to acquire Fe using specialized Fe-chelating compounds (*e.g.,* siderophores) and several Fe transport systems ^18,19^. Species-specific associations between bacteria and algae may arise due to exchange of Fe-siderophores ^20^, but a molecular level understanding of how Fe availability alters algal dissolved organic matter (DOM) production and subsequent processing by heterotrophic microbial taxa has not been established. Fe limitation can lead to the colimitation of carbon and Fe for the bacterial community, impacting Fe dependent carbon metabolism within cells ^5,21,22^.

There is ample evidence that molecular composition of algal DOM is an important driver of microbial metabolic activity and microbial community assembly, though the impact of Fe on these processes is unknown. Activity-based approaches quantifying carbon flow have demonstrated that both quantity and quality of algal DOM exert selective pressure on microbial communities ^23,24^ and reveal functional guilds of bacteria consuming and remineralizing algal DOM ^25^. Current mass spectrometry approaches have led to the identification of specific DOM substrates that may drive bacterial community assembly. For example, bacterial taxa associated with the diatom *Thalassiosira pseudonana* take up distinct classes of exuded organic molecules ^26^.

Additionally, the model diatom *Phaeodactylum tricornutum* exudes an organic acid, 4-hydroxybenzoic acid, which promotes the growth of specific bacterial taxa ^27^, and the coccolithophore *Geocapsa huxleyii* secretes a similar compound, benzoate, which also promotes specific bacterial taxa ^28^, indicating common molecular mechanisms of selectivity across highly phylogenetically divergent algal groups. *P. tricornutum* also exudes volatile organic carbon (VOC) compounds, such as benzene, which are consumed by specific bacterial taxa, highlighting this DOM class as potentially important as selective substrates ^29^. These studies demonstrate the importance of obtaining molecular-level resolution, ideally paired with activity-based measurements of carbon flow, to predict shifts in cycling of algal-derived organic matter in response to Fe limitation.

In this study, we tested the hypothesis that shifts in host phototroph metabolism and health, driven by Fe limitation, impact microbiome dynamics via exudation of specific DOM exometabolites. We employed a model diatom host, *P. tricornutum*, whose response to Fe limitation has been extensively characterized ^16,30–32^, and a *P. tricornutum* microbial enrichment culture that depends on the diatom-derived DOM for growth^27,33,34^. Using this experimental system grown under Fe-replete and Fe-deplete condition, we measured the microbial response to Fe-limited *P. tricornutum*. We first used nanometer-scale secondary ion mass spectrometry (nanoSIMS) to measure single-cell diatom and bacterial Fe:C ratios, and then combined nanoSIMS with stable isotope probing (nanoSIP) to measure bacterial incorporation of diatom exudate and activity (via ^13^C-bicarbonate additions and ^15^N-ammonium additions, respectively). To identify potential substrates that could be linked to the single-cell activity, we used ultra-high-resolution liquid chromatography (LC)-MS enabled exometabolomics ^35,36^ on the community and axenic (diatom-only) cultures under the different Fe conditions. We then compared community composition between the two Fe conditions to identify the metabolic potential of the taxa in the Fe-limited diatom microbiome which may be driving differential exometabolite consumption. Together, these results provide metabolomic and genomic information to further a predictive understanding of the role of Fe availability in regulating how microbial communities process diatom-derived organic carbon in the marine ecosystem. Additionally, as *P. tricornutum* is a model biofuel diatom, this work provides industrial operators with a potential lever in the form of Fe availability to alter microbiome feedbacks to host photosynthetic processes and productivity in engineered systems.

## Methods

### Organisms and culturing

The axenic diatom, *Phaeodactylum tricornutum* Bohlin strain CCMP 2561, and a phycosphere enrichment culture containing *P. tricornutum* CCMP 2561 (hereafter “enrichment culture”) with a mixed community of heterotrophic bacteria ^34^ were used for a series of Fe-limitation experiments. Axenicity was confirmed prior to the experiments using microscopy. Experiments were conducted using enriched artificial saltwater (ESAW; ^37,38^) without sodium metasilicate, and a sterile stock of 0.01 M FeCl_3_ 6H_2_O + Na_2_EDTA 2H_2_O was prepared to bring the final media to either 2 µM FeCl_3_ (Fe-replete) or 10 nM FeCl_3_ (Fe-deplete). The concentration of total Fe, measured by inductively coupled plasma mass spectrometry (ICP-MS, Agilent 7850), was 2.69 µM and 10.09 nM in the Fe-replete and Fe-deplete media, respectively. Unless specified, all experiments were carried out in ∼250 mL volume in 500 mL sterile polycarbonate Erlenmeyer flasks (screw cap, vented) previously soaked in 10% hydrochloric acid (HCl) for a minimum of 48 h, rinsed 6x with MilliQ water and sterilized by autoclaving. Media was prepared in sterile, HCl-washed polycarbonate bottles, and plastic utensils were used to measure media components.

All cultures were grown in batch mode with shaking (90 rpm, 22°C, 12h:12h light:dark, 3500 lux illumination), and growth was monitored using chlorophyll *a* fluorescence (Trilogy Fluorometer, Turner Designs, San Jose, CA, USA, Chlorophyll *In Vivo* Module) and flow cytometry (Thermo Attune NxT). Detail on the flow cytometry protocol is described in Supplementary Methods S1.

For all experiments, the cultures were pre-acclimated by culturing the parent inoculum in ESAW containing no added Fe for two transfers before the start of the experiment to deplete intracellular Fe. To assess photosystem health a Mini-PAM-II (Heinz Walz GmbH, Effelitrich, Germany) was used, 1 mL samples were acclimated for 15 minutes in the dark and then exposed to a white saturating light pulse prior to determining maximum theoretical photochemical quantum yield of photosystem II (Fv/Fm). To prepare samples for nanoSIMS Fe:C measurements, samples were fixed with 3.4% 0.2 µm filtered formaldehyde and stored at 4°C in the dark for <1 week, after which they were filtered onto 0.2 µm polycarbonate filters (MilliporeSigma, Burlington, MA, USA), washed 3x with 0.2 µm filtered MilliQ water and dried. Portions of filters were cut out and adhered to aluminum disks using conductive tabs and sputter coated with ∼5nm of gold, as described previously ^25^.

### Nanoscale stable isotope probing (nanoSIP) experiment

To quantify the transfer of *P. tricornutum* derived C into the microbiome under Fe-limitation, an experiment was performed using stable isotope probing and nanoSIMS imaging (nanoSIP). Here, the axenic and enrichment *P. tricornutum* cultures were acclimated and grown in Fe-deplete and Fe-replete media in triplicate (“core incubation”). After four days of growth, each flask was subsampled for nanoSIP incubations. For labeled treatments, 10 mL from each flask was spiked with 500 µM ^13^C-sodium bicarbonate (99 atom% ^13^C; Cambridge Isotopes Laboratories, Cambridge, Massachusetts) and 100 nM ^15^N-ammonium chloride (99 atom % ^15^N; Cambridge Isotopes Laboratories, Cambridge, Massachusetts) and split into triplicate acid-washed 5 mL round-bottom Hungate tubes. For bacteria-only controls, 10 mL subsamples were first filtered through a 0.8 µm syringe Supor filter (Pall, Port Washington, NY) to remove diatom cells and then label added. Killed controls were prepared by fixing 5 mL from each flask to 3.4% 0.2 µm filtered formaldehyde for at least 15 minutes in the dark, after which the label was added and incubated in the dark. For unlabeled controls, 5 mL from each flask was spiked with 500 µM sodium bicarbonate and 100 nM ammonium chloride, both natural-abundance. All tubes were incubated for 24 h under the same conditions as the core incubation (except for the killed control), fixed with 3.4% 0.2 µm filtered formaldehyde, and processed as described above for the nanoSIMS filter preparation. Samples for flow cytometry were collected at the start and end of the nanoSIP experiment by fixing a 200 µL aliquot of the sample with glutaraldehyde (0.25%).

### Nanoscale secondary ion mass spectrometry (nanoSIMS) and nanoSIP analysis

Single-cell relative Fe:C ratios (^56^Fe^+^/^12^C^+^) were generated by analyzing the filtered samples using a NanoSIMS 50 (CAMECA) at Lawrence Livermore National Laboratory equipped with a Hyperion II inductively coupled RF plasma ion source (Oregon Physics). Analyses were conducted with a focused 200 pA O-beam scanned over a 30 µm x 30 µm square area and quantitative secondary ion images were collected for ^12^C^+^, ^31^P^+^, ^40^Ca^+^, ^56^Fe^+^, and ^63^Cu^+^ on individual electron multipliers in pulse counting mode generating images with 256 x 256 pixels. Metal ion peaks were identified using NBS610 glass (NIST). Each area was scanned 50 times (cycles) with 1 ms pixel-1 dwell times; ion count rates stabilized after the first 5 cycles, and therefore the first 5 cycles were excluded. Ion image data were processed using L’Image software (L.Nittler, Carnegie Institution of Washington) run on IDL (Harris Software). Cells were identified using the ^40^Ca^+^ ion images, regions of interest (ROIs) manually drawn around diatom and bacterial cells, and ^56^Fe^+^/^12^C^+^ ratio averages and standard error of the mean of the cycles were calculated and exported for each ROI (Supplementary Table 1).

The nanoSIP imaging analyses to generate ^13^C and ^15^N single cell enrichment values was performed with a prototype NanoSIMS (CAMECA) at Lawrence Livermore National Laboratory based on the NanoSIMS-HR physics and electronics ^39^ but equipped with a high-brightness Cs^+^ source (“low temperature ion source”, or LoTIS)^40^. Secondary electron images and quantitative secondary ion images were simultaneously collected for ^12^C_2_ ^-^, ^13^C^12^C^-^, ^12^C^14^N^-^, and ^12^C^15^N^-^ on individual electron multipliers in pulse counting mode. Target areas were sputtered to a depth of ∼60 nm with an initial beam current of 150 pA to achieve sputtering equilibrium. A 1 pA Cs^+^ was rastered over 15 µm x 15 µm analysis areas with a dwell time of 0.5 ms pixel^-1^ for 15 scans (cycles) and generated images containing 512 x 512 pixels. ROIs were manually drawn in L’Image (http://limagesoftware.net) around diatom cells (avoiding attached bacteria), attached bacteria (cells associated with a diatom cell) and free-living bacterial cells using the ^12^C^14^N^-^ ion images. The isotopic ratio averages and standard error of the mean of the cycles were exported for each ROI (Supplementary Table 2). We calculated *X_net%_*, which represents the percent of a cell’s biomass that originated from the isotope-labeled substrate, using the single cell measurements of isotopic enrichment (f*_Xf_*) and taking into account the level of isotope labeling in the substrate (f*_X_*_s_) and the isotope labeling of the biomass at the start of the incubation (f*_X_*_i_, killed controls)^41^.

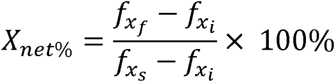

For bacterial the bacteria were consuming diatom-derived exuded organic carbon substrate (f*_Xs_*) of unknown C atom %, so we used average diatom cell C APE for each treatment, with the assumption that diatom-exuded carbon would be at a similar ^13^C APE as the diatom cells in that same sample. Given this assumption we term the bacterial incorporation of new C as *C_net%_^’^*, to distinguish it from the other *X_net_*_%_ calculations, which can be calculated more precisely.

### Exometabolomics and microbiome response to Fe limitation experiment

The diatom exudate was profiled using exometabolomics, and the microbiome response was monitored using partial 16S rRNA gene sequencing. The parent inoculants were grown in Fe-replete ESAW for several generations to achieve acclimation. Following the acclimation period, axenic *P. tricornutum* and the enrichment culture were inoculated in triplicate into Fe-replete or Fe-deplete media to obtain similar starting fluorescence across all flasks (1639 ± 49 relative fluorescence units). After 5 days and 9 days of growth, the cultures were filtered onto sterile 0.2 µm Supor membrane filters (Pall, Port Washington, NY). The filter was immediately frozen at −80°C for DNA extraction, and the filtrate was collected directly into 50 mL Falcon tubes and frozen at −80°C for exometabolomics. Media blanks were collected by filtering the Fe-replete and Fe-deplete ESAW media used for the experiment through a 0.2 µm Supor membrane filter, collecting the filtrate into 50 mL Falcon tubes and storing at −80 °C.

### Bulk dissolved organic carbon (DOC)

In order to measure DOC in the exudate under Fe replete and Fe deplete conditions, the enrichment culture was grown in the same manner as described above. After 5 and 9 days of growth, 25 mL of exudate was filtered through Supor filters pre-rinsed with sterile MilliQ water, placed into pre-combusted (450°C for >4 h) 40 mL TOC vials, acidified by adding 2µL 4M HCl / ml of sample, and stored at room temperature until analysis. DOC was measured using a Shimadzu TOC TOC-V analyzer by the Carlson DOM analysis lab at the University of California Santa Barbara, according to protocols described in Halewood et al ^42^.

### LC-MS and exometabolomics annotation

To prepare for metabolomic analysis, samples were solid phase extracted following standard methods^35,43,44^ which are described in Supplementary Methods S1. Cyanocobalamin (Sigma-Aldrich) was used as an internal standard. To ensure reproducibility over the course of analysis, a pooled sample was generated by combining equal volumes of each sample and was analyzed several times during LC-MS analysis. Although specific extraction methods cannot capture all pools of DOM, the PPL column used have been shown to recover 40-60% of DOM from marine samples^45^. Highly polar molecules are not retained and missed by this analysis.

LC-MS analysis was conducted with an ultrahigh performance liquid chromatography system (Thermo Vanquish Horizon) coupled to an orbitrap mass spectrometer (Thermo Orbitrap IQ-X). The specific details of the LCMS run are in Supplementary Methods S1. The standard deviation of the cyanocobalamin internal standard peak area was 15%, and the retention time variability was 0.01 min across the data set. One sample (axenic replicate grown in 10nM Fe) was excluded from further analysis due to low LCMS signal intensity.

After data acquisition, a metabolome feature list containing the measured *m/z*, retention time, and abundance of detected metabolites across the samples was generated with MS Dial ^46^. Peak detection was performed with a minimum peak height of 5e^4^ and mass slice width of 0.05 Da. Features were annotated based on MS/MS fragmentation matching to the MSDial ESI(+)-MS/MS reference library from authentic standards (VS17). Detail on MS/MS matching and feature alignment across samples is described in Supplementary Methods S1. For blank filtering, the sample max/blank average was set to 5 fold change. This yielded a final list of 21,378 features, with 18,771 features after blank filtering. Overall, 615 features were annotated by MS/MS matching (3% of blank-filtered features). Fragmentation spectra for features of interest reported in the main figures were also manually inspected to ensure confident assignment.

Molecular formula were assigned using CoreMS as described previously ^47,48^. Mores detail on the formula assignment are provided in Supplementary Methods S1. Assignments were made up to a maximum of 400 *m/z*, above which formula assignments based on Orbitrap MS became less reliable. Assigned formulas were combined into a comprehensive formula library list by tabulating each assigned formula in a unique retention time window, consolidating features with overlapping m/z, with selected assignments based on maximum confidence score, and filtering out features with anomalous m/z error beyond the 99.99% confidence interval. Assignments were annotated with stoichiometric classifications based on the multidimensional criteria of Rivas-Ubach et al ^49^. This molecular formula library was used to annotate features from the MSDial feature list following the approach of Coffey et al.^35^ using an *m/z* tolerance of 10 ppm. Isotopologues were excluded from formula assignment.

### Microbial community analysis using 16S rRNA amplicon and metagenomic sequencing

Samples for 16S rRNA amplicon sequencing were generated from the exometabolomics experiment (see above) by extracting whole DNA using a phenol chloroform extraction procedure ^50^ (details in Supplementary Methods SI), amplified using Illumina V3-V4 universal primers^51^, and sequenced on the Illumina MiSeq platform (Laragen Inc., Culver City, CA). Metagenome assembled genomes (MAGs) were generated from a representative Fe-limited *P. tricornutum* enrichment community that was filtered onto a 0.2 µm Supor filter and extracted as described for the 16S rRNA amplicon library. The metagenome was sequenced through the Joint Genome Institute (JGI) using the Illumina NovaSeq S4. Details on 16S rRNA amplicon library and metagenome generation, bioinformatic processing, and statistical analyses are described in Supplementary Methods S1.

### Isolation of a low Fe adapted bacterial taxa in the enrichment community

We attempted to isolate taxa that were abundant in the Fe-limited enrichment culture by plating the low Fe acclimated enrichment culture onto a variety of media (ESAW, 100% ZoBell high nutrient media, 20% ZoBell, sterilized exudate from the Fe-limited culture) and incubating in the dark at 22°C. Whole genome sequencing of colonies of interest was done via Genewiz from Azenta Life Sciences (South Plainfield, NJ). Here, DNA from the colonies was extracted using the phenol:chloroform method described above and total DNA was sequenced 2×150 bp using the Genewiz Whole Genome Sequencing NGS method. Kbase was used to quality control, assemble, and annotate genomes. Paired-end reads were uploaded and trimmed using Trimmomatic v0.39, and read quality was assessed using FastQC 0.12.1. SPAdes v 3.15.3 was used to assemble using default parameters, and RASTtk v1.073 was used for gene calling and annotation.

### Fe-limitation experiment of low-Fe adapted Roseibium C24 strain

We were able to isolate a strain of *Roseibium aggregatum* sp. (strain “C24”). To test the response of C24 to Fe limitation in isolation, six baffled flasks were prepared with 25mL each of Enriched Artificial Seawater Medium (ESAW) supplemented with 5 mM glucose as the sole carbon source. Three flasks were prepared for each Fe replete and deplete condition, supplemented with sterilized 2 µM and 10 nM FeCl_3_, respectively. For inoculation, C24 was grown to saturation in ESAW, 5 mM glucose, iron replete media, then washed twice with PBS before inoculating the flasks. Cultures were incubated at 22°C with 250 RPM shaking for 48 hours and sampled periodically for ICP-MS and flow cytometry. Collection for ICP-MS involved centrifuging 1 mL of bacterial culture at 21300 x g for 1 min at room temperature, supernatant removed and pellet stored at –20°C. 100 µl samples were collected for flow cytometry, fixed with glutaraldehyde (final concentration 0.25%), and stored at –80°C. Cells were prepared for ICP-MS by resuspending pellets in 1 mL of 5_% nitric acid in a 96 deep well plate, and then metals were quantified using an Agilent 7850 ICP-MS instrument (Agilent Technologies) equipped with a microplate adaptor. See Supplementary Methods S1 for details on ICP-MS analysis.

### Statistical analyses

To compare the nanoSIMS isotope data, the Mann-Whitney test was used to calculate statistical significance between the Fe-replete and Fe-deplete condition in GraphPad Prism 10. The ICP-MS results for the *Roseibium* experiment were compared between the Fe-replete and deplete condition using an unpaired t-test in GraphPad Prism 10. Metaboanalyst 6.0 ^52^ single-factor analysis was used for exometabolomics comparisons between Fe-replete and Fe-deplete conditions, or between the axenic culture and the enrichment culture. Here, peak intensity values were normalized by the sum of that metabolite across samples and log_10_ transformed. Fold change and significance (unpaired t-test) was calculated for each exometabolite, comparing the Fe-replete to the Fe-deplete condition or axenic to enrichment culture. For comparing exometabolites differences between the Fe-treatments, we focused on characterizing the exometabolites that were significantly increased in the Fe-deplete condition because an increased diatom cell abundance in the Fe-replete cultures may cause an increased abundance of an exometabolites. Exometabolites that had positive log_2_-fold changes in the enrichment culture samples relative to axenic samples were considered “produced”, and those with negative log_2_-fold changes were considered “consumed”. Produced exometabolites could have been exuded from the bacteria or were exuded by the algae when the microbiome was present. Consumed exometabolites could have occurred via bacterial uptake for assimilation of dissimilatory processes, or the exometabolite was extracellularly transformed. Within R, the VEGAN package v.2.6-4 ^53^ was used to construct nonmetric multidimensional scaling (NMDS) plots using Bray-Curtis distances calculated using the normalized exometabolite abundances. More detail on statistical analyses is available in Supplementary Methods SI.

### Data availability

The raw metagenome data is available through JGI under Project ID 1495622. Raw amplicon data can be downloaded via Zenodo (https://zenodo.org/records/18944121). Raw MS data is available in the MassIVE repository (https://massive.ucsd.edu/) under accession number MSV000101105. Code for figures and analyses are available through GitHub (https://github.com/negilbert/PtFeExo_code).

## Results

### Growth and physiological responses of P. tricornutum and bacteria to Fe availability

Fe availability significantly affected diatom and bacterial abundances and growth. The growth rates of *P. tricornutum* in the microbial enrichment culture were 0.19 d^-1^ and 0.09 d^-1^ under Fe-replete and Fe-deplete conditions, respectively (Fig. 1A, Supplementary Table 3). Bacterial growth in the enrichment culture was also significantly hampered under Fe-deplete conditions (Fig. 1B). Bacterial growth rates during exponential phases were 1.07 d^-1^ and 0.35 d^-1^ for Fe-replete and Fe-deplete conditions, respectively (Fig. 1, Supplementary Table 1). By day 9, bacterial abundance was ∼20-fold higher in the Fe-replete cultures, reaching up to 1.7 x 10^7^ cells/mL (Fig. 1B). Fe-deplete bacterial abundances reached a maximum of 8.05 x 10^5^ cells/mL at day 9 (Fig. 1B). In the Fe-deplete cultures, there was a steady decline in the ratio of bacteria to diatom cells from ∼1 to ∼0.37 bacterial cells per diatom cell, indicating that bacteria had a slower growth rate than the diatom in response to Fe limitation d (Supplementary Fig. S1).

**Figure 1.**
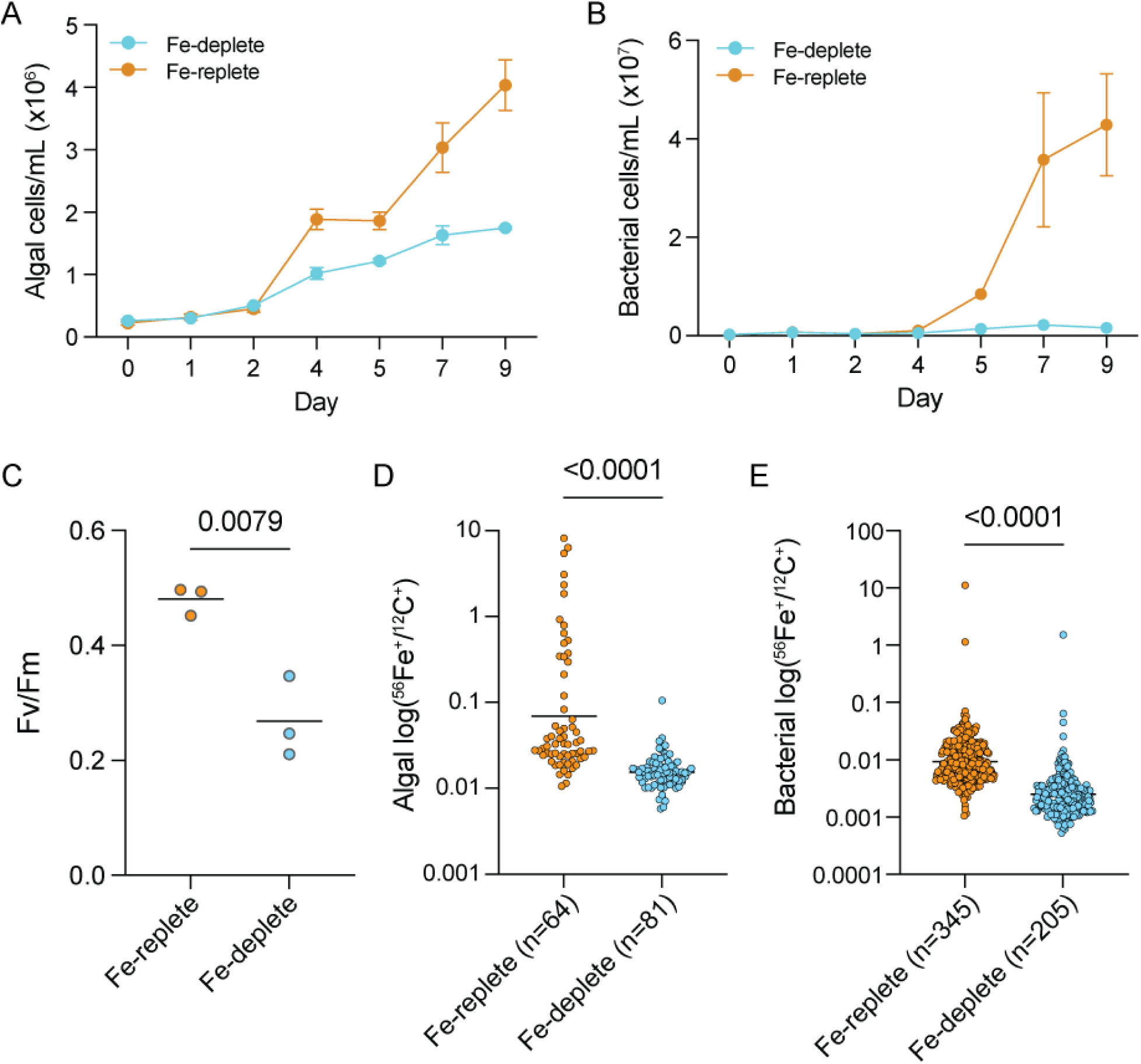
Growth and physiological response of the *P. tricornutum* enrichment community to Fe-limitation. **A)** Algal growth under Fe-replete (2 µM-added Fe) and Fe-deplete (10 nM-added Fe) conditions. B) Bacterial growth of the *P. tricornutum* microbiome enrichment culture. **C)** Photosynthetic efficiency measured as Fv/Fm for the *P. tricornutum microbiome* enrichment culture. P-value calculated using an unpaired t-test. NanoSIMS derived Fe (^56^Fe^+^) to carbon (^12^C^+^) ratios in single cells at day 9 for **D)** Algae and **E)** Bacteria. P-values calculated using the Mann-Whitney test. The numbers (n) of cells analyzed are shown in the x-axis.

There were also significant differences in the physiological responses of the diatom and bacteria in the enrichment cultures between the treatments. *P. tricornutum* photosynthetic efficiency of PSII was significantly (p=0.0079) reduced in the Fe-deplete cultures (Fv/Fm = 0.27 ± 0.07) compared to the Fe-replete cultures (0.48 ± 0.03, Figure 1C), a common symptom of Fe deficiency stress. The single-cell ^56^Fe:^12^C ratio, derived from nanoSIMS analysis of the day 9 samples, was also significantly reduced (p<0.0001) for *P. tricornutum* and bacteria (Fig. 1D & E), indicating a reduction in biomass Fe concentrations for both the diatom and the bacterial community under Fe-limitation.

### Bacterial incorporation of *P. tricornutum*-derived carbon under Fe limitation

We aimed to quantify diatom C-fixation and bacterial incorporation of C as a function of Fe availability using nanoSIP (Fig. 2). To do so, we sub-cultured enrichment cultures grown for 4 days in the Fe-deplete versus Fe-replete media, then spiked with ^13^C-bicarbonate and trace amounts of ^15^N-ammonium to track C incorporation compared to activity over 24h. Since the media was not removed, there was carryover unlabeled “old” *P. tricornutum*-derived DOC also present at the start of the labeling. We calculated net incorporation of newly fixed carbon into diatom biomass based on ^13^C enrichment (C*_net_*_%_) ^41,54^. *P. tricornutum* single-cell C-fixation was significantly reduced (p<0.0001) in the Fe-deplete condition, with a mean C*_net%_* of 3.93% compared to 7.10% per day in the Fe-replete condition (Fig. 2C, Fig. S2, Supplementary Table 5).

**Figure 2.**
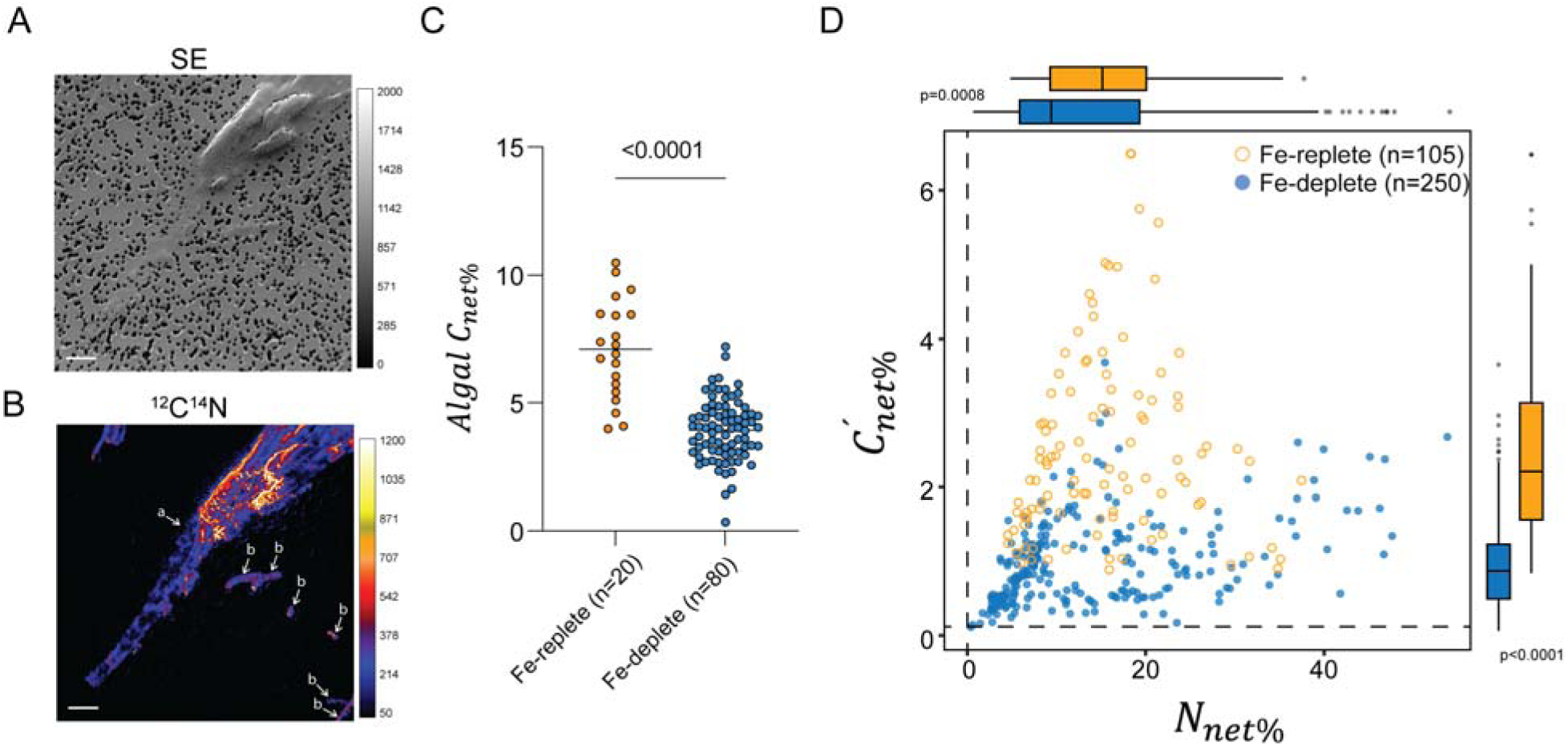
Stable isotope probing and paired single-cell analysis (NanoSIMS) demonstrate how the production and flow of diatom-derived carbon in the *P. tricornutum* enrichment community shifts with Fe-availability. Representative NanoSIMS images showing **A)** Secondary electron (SE) counts and **B)** ^12^C^14^N^-^ counts used to draw algal “a” and bacterial “b” regions of interest. **C)** Algal carbon fixation, measured as net incorporation of ^13^C-bicarbonate into single *P. tricornutum* cells (C*_net%_*). The lines indicate the mean C*_net%_* values. **D)** Single-celled bacterial incorporation of algal organic carbon versus activity as measured by net incorporation of ^15^N-ammonium (*N_net%_*). Bacterial *C_net%_^’^* was calculated for each Fe-condition using the average ^13^C enrichment of the diatom (see Methods). Dashed lines indicate the average background *C_net%_^’^* or *N_net%_* calculated for killed and bacteria-only controls. Open orange circles indicate Fe-replete cells and closed blue circles indicate cells in the Fe-deplete condition. The marginal boxplots indicate the means and spread of *C_net%_^’^* and *N_net%_* for each Fe-condition. The p-values comparing *C_net%_^’^* or *N_net%_* between iron-deplete and replete conditions were calculated using the Mann-Whitney test.

We used diatom ^13^C enrichment under the two conditions to estimate bacterial *C_net%_^’^*, which is a measure of the percentage of total bacterial biomass that was newly synthesized from the diatom-derived C in each bacterial cell, in the Fe-deplete versus Fe-replete conditions. We found that bacterial *C_net%_^’^* was significantly (p<0.0001) reduced in the Fe-deplete cultures (n=250 cells, 0.997%) compared to the Fe-replete cultures (n=105 cells, 2.491%; Fig. 2D, Supplementary Fig. S2, Supplementary Table 5). By contrast, we found that the average activity of the Fe-deplete bacterial cells, as measured by net incorporation of ^15^N-ammonium (*N_net%_*, Fig. 2D) was only 10% lower than in the Fe-replete cultures (13.96% vs. 15.52%, respectively, p=0.0008), and there was a population of Fe-deplete cells that had higher *N_net%_* values (some >40%) than those in the Fe-replete cultures (Fig. 2D, Supplementary Table 5). Furthermore, by comparing *C_net%_^’^* against *N_net%_*, we found that cells had higher incorporation of ammonium N per unit diatom C (Fig. 2D), suggesting lower incorporation of diatom-derived C despite high activity. The *N_net%_* coefficient of dispersion (CD) for the Fe-deplete community was 54.92% compared to 37.92% in the Fe-replete community (Supplementary Table 5), indicating that under Fe limitation there was a greater degree of single cell heterogeneity, which could stem from either a more diverse set of active taxa or a shift in metabolic activity of the same active taxa. Further, we found a significantly (p<0.0001; Hartigans dip test) bi-modal trend for the Fe-deplete community when comparing *C_net%_^’^*: *N_net%_* (Supplementary Fig. S2E), indicating a population of cells with distinct metabolic activity in the Fe-deplete community relative to the Fe-replete condition. From assessment of enrichment of bacterial cells that were visibly attached to diatom cells compared to unattached bacterial cells, there was no significant difference in ^13^C or ^15^N enrichment under either replete or deplete conditions (Supplementary Fig. S2D). Given that the diatom cell counts did not show cell lysis on day 4 in the Fe-deplete condition (Fig. 1A), this indicates that the distinct bacterial metabolic activity observed in the Fe-deplete condition is unlikely to be the result of consumption of dead or lysed diatoms. Altogether, the data suggest that this distinct population of bacteria in the Fe-deplete condition were consuming more of the unlabeled carryover DOC.

### Composition of diatom exudate changes with Fe availability over time

The nanoSIP results indicated that active populations under Fe-limitation consumed more low-quality resources than under Fe-replete conditions, so we characterized how the diatom exudate shifted using exometabolomic profiling of the axenic *P. tricornutum* culture. Previous studies have demonstrated that the exudate of *P. tricornutum* significantly differs from the cellular pool ^55^, and we did not observe evidence of significant cell lysis/death in Fe-limited conditions. Thus, we interpret the exometabolites as actively and/or passively exuded. The results revealed that Fe-limitation drastically altered exometabolite pools of axenic *P. tricornutum* (Fig. 3). These patterns were time-dependent, where day 5 exometabolomic profiles differed from day 9 exometabolomic profiles, although there was a greater separation between Fe condition at day 9 (Supplementary Fig. S4A). At day 9, there were 1308 (530 had log_2_(fold change) ≥ 2) exometabolites that had significantly (p<0.05) higher abundance under Fe limitation, and 3086 (2974 had log_2_(fold change) ≥ 2) that had significantly lower abundance under Fe limitation (Supplementary Table 7). Exometabolites more abundant in the Fe-deplete conditions included lipids, amino acids/peptides, and heterocyclic aromatics. Using the traditional annotation approach that relies on public databases (*e.g.,* MSDial), we were able to annotate only 13 exometabolites that had confident MS/MS matches (via manual curation) to a reference (Fig. 3A). These included vitamin degradation products, nucleotides/nucleoside-like molecules, amino acids, lipids, and putative signaling/facilitator molecules and antimicrobials (Supplementary Table 7). Then, using CoreMS annotation for confident molecular formula assignments (4,798) of the unannotated features, we found that the stoichiometric composition of the exometabolites was distinct between Fe-replete and Fe-deplete exudate (Fig. 3B). Across all features that were significantly enriched in the Fe-deplete exudate (1,308), 755 were assigned a molecular formula using CoreMS (Supplementary Table 7). Based on classes predicted from the stoichiometric composition, the Fe-deplete exudate was predominantly composed of lipids and phytochemicals (Fig. 3B). The Fe-deplete exudate had significantly (p<0.05) higher mean average retention time (min) and oxygen:carbon ratio (Supplementary Fig. S3).

**Figure 3.**
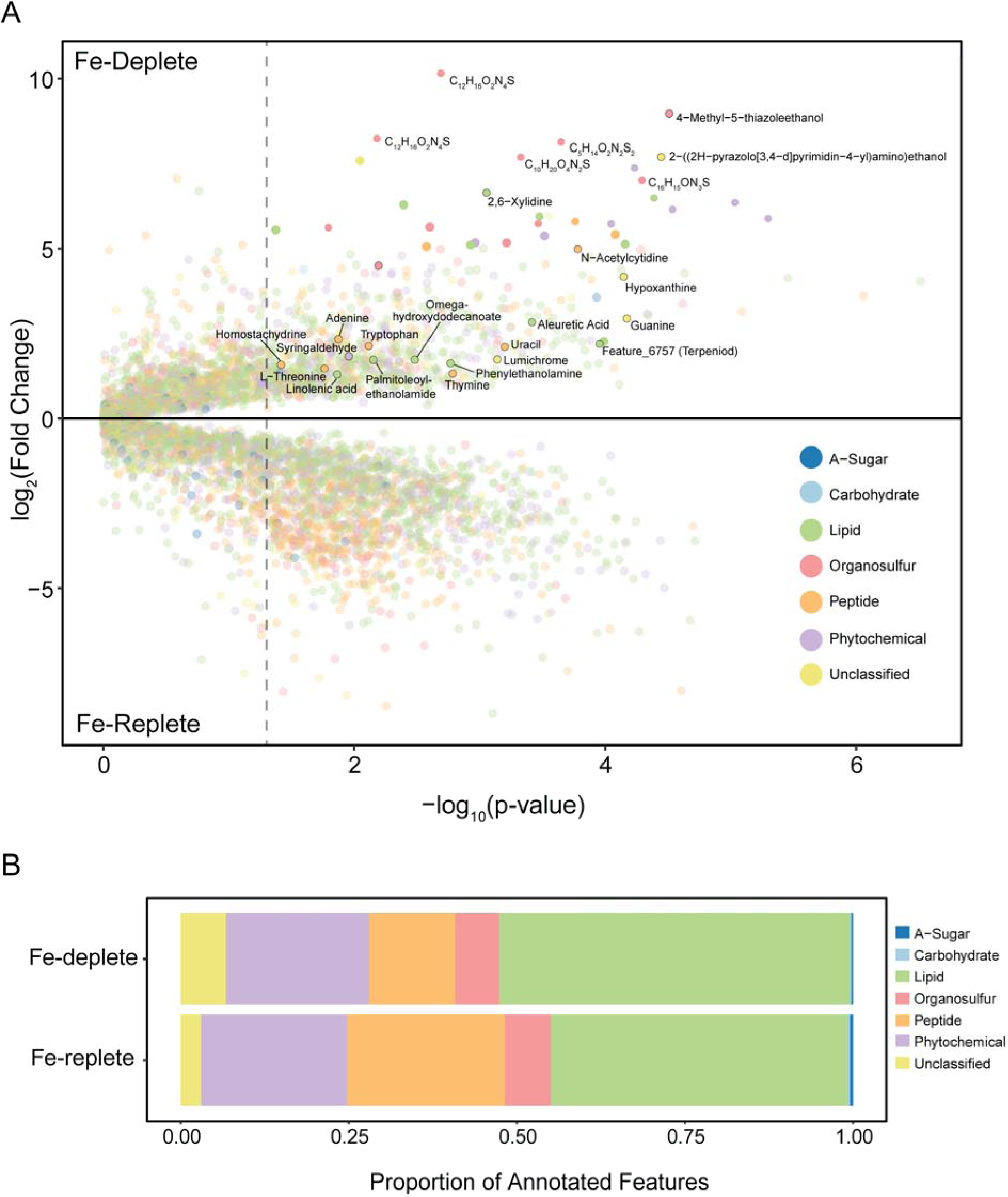
*P. tricornutum* exudate (exometabolites) composition shifts with Fe-availability. (A) Fold change analysis of annotated exometabolites in the axenic *P. tricornutum* exudate, between the Fe-replete and Fe-deplete axenic cultures at day 9. P-value calculated using unpaired t-test for each exometabolites. The dashed line indicates a p-value of 0.05. Colors represent stoichiometric group for metabolites annotated using the CoreMS pipeline. Unclassified means the CoreMS molecular formula did not fall within the 7 stoichiometric groups. Statistically significant exometabolites that were enriched in the Fe-deplete condition are further annotated. Exometabolites annotated using MSDial are shown with the metabolite name. Exometabolites annotated using CoreMS but not using MSDial are opaque if they had a log2(FC) >=5, Molecular formulae is shown for exometabolites with the highest log2 fold change values. Feature_6757 (Terpeniod) = (1S,4R,5R,9S,10R,13R)-5,9-dimethyl-14-methylidenetetracyclo(11.2.1.01,10.04,9)hexadecane-5-carboxylic acid. (B) Stoichiometric summary of significantly increased exometabolites in the Fe-deplete or Fe-replete conditions. Undetected/not annotated exometabolites using the CoreMS pipeline are not shown.

### Bacterial consumption and production of exometabolites in the diatom enrichment community

We next examined shifts in the exometabolome pool when the microbiome was present. By comparing exometabolite abundance between the enrichment culture and the axenic *P. tricornutum* exudate, we inferred whether an exometabolite was consumed (depleted in the enrichment culture) or produced (accumulated in the enrichment culture) when the microbiome was present. There were 193 compounds that were significantly consumed in the Fe-deplete condition (Fig. 4). From the k-means clustering of log_2_-fold change values between the enrichment and axenic cultures in Fe-replete or Fe-deplete conditions at day 9, we saw these fell into a distinct k-means cluster (Supplementary Fig. S4, Supplementary Table 9). Structurally annotated metabolites that exhibited greater consumption under Fe-deplete conditions than Fe-replete conditions included the purine guanine, pyrimidines thymine and uracil, the nucleoside deoxyadenosine (log_2_fc = 6.76, p=0.0007), the amino acid tryptophan, and the aromatic compounds 2,6,-xylidine, syringaldehyde, and benzylideneacetone (Fig. 4B, Supplementary Table 9). Several other amino acids, peptides, nucleotides and nucleosides-like exometabolites were also significantly consumed under both Fe-deplete and replete cultures, including hypoxanthine, uracil, adenosine, 5’-methylthioadenosine, adenine, guanine, N-acetylcytidine and thymine (Fig. 4B, Supplementary Table 9). Overall, a distinct set of exometabolites was consumed uniquely or had higher consumption under Fe-deplete conditions.

**Figure 4.**
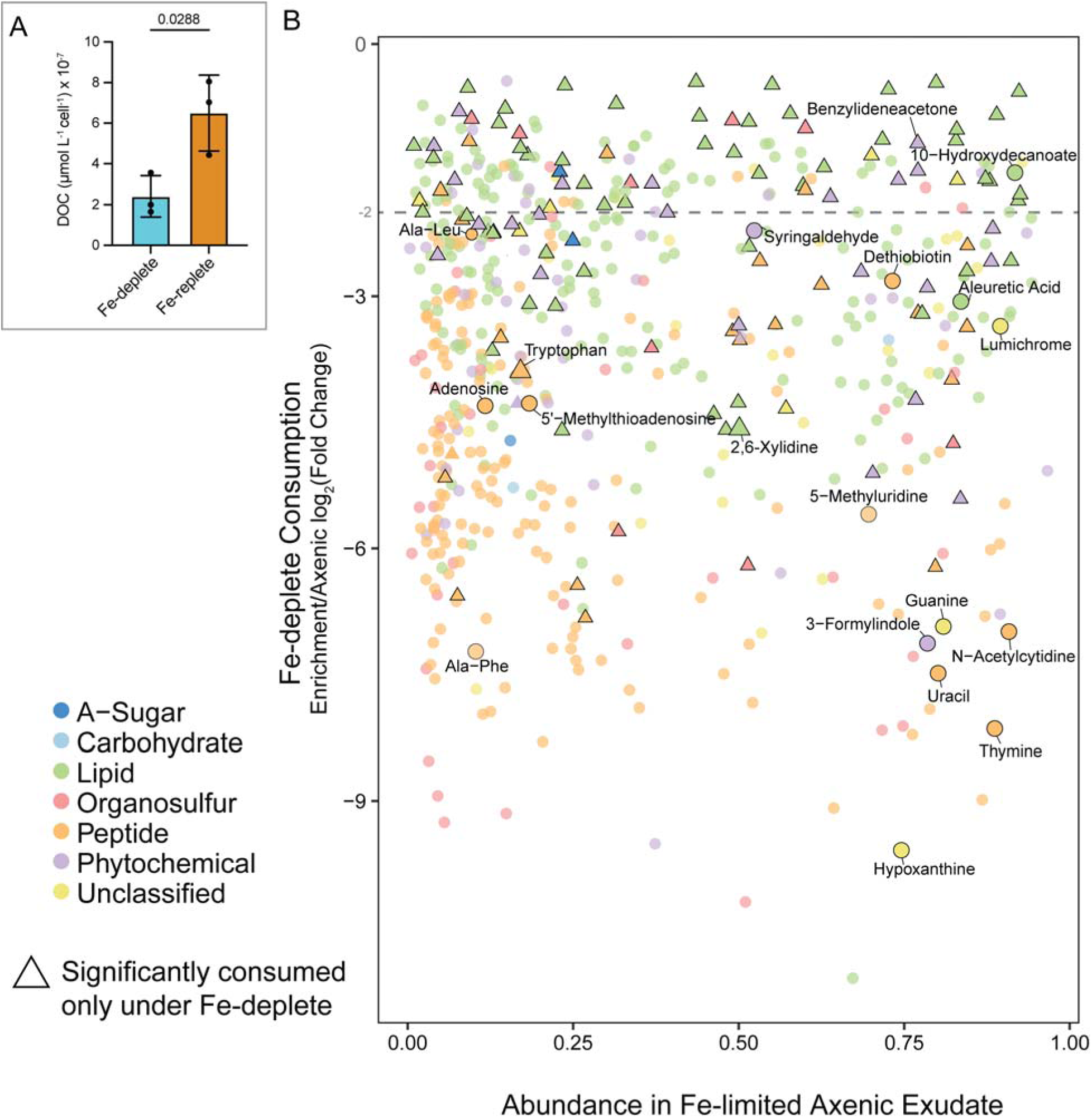
Consumption of exometabolites in the Fe-limited microbiome relative to availability in Fe-limited exudate. A) Total dissolved organic carbon (DOC, µmol per L per algal cell) in the enrichment community at day 9. p-value calculated using an unpaired t-test. B) Consumption of exometabolites in the microbiome enrichment community shown via log_2_(Fold Change) values between the enrichment vs the axenic *P. tricornutum* culture in the Fe-deplete condition. Only significantly (p<0.05, unpaired t-test) different exometabolites in the Fe-deplete condition are shown. Triangles show exometabolites only significantly consumed in the Fe-deplete condition. Circles show exometabolites significantly consumed in both Fe-deplete and Fe-replete conditions. Exometabolite abundance in the Fe-deplete axenic exudate was normalized by the sum peak intensity across both Fe-deplete and Fe-replete samples.

We also measured total DOC in the diatom exudate of the enrichment cultures at day 5 and 9 of growth. By day 9, the concentration of DOC normalized to diatom abundance was reduced 3.5 fold (p=0.0288) in the Fe-deplete cultures (0.0002 ± 0.0001 µmol L^-1^ cell^-1^) compared to the Fe-replete cultures (0.0007 ± 0.0002 µmol L^-1^ cell^-1^, Fig. 4A & Supplementary Table 6), indicating decreased exudation and/or increased bacterial consumption under Fe-limitation. The DOC: diatom cell abundance was highly variable in the day 5 samples (Supplementary Fig. S5A), so no direct comparison could be made with the nanoSIP data. Nevertheless, the nanoSIP data from day 5 indicated less diatom-derived carbon was incorporated on average into single bacterial cells relative to the Fe-replete condition (Fig. 2D) supporting decreased exudation. We calculated net bacterial biomass production (C*_net%_*-_total_) for individual growth experiments (number of bacteria x mass per cell x C*_net%_*) ^56^, and found that there was 3.75-fold less diatom-derived carbon incorporated into bacterial biomass under Fe-limitation (Supplementary Fig. S5B). Although this does not take into account bacterial respiration, together these results suggest that total diatom organic carbon exudation decreased under Fe limitation, along with less flow of this carbon into bacterial biomass.

### Microbial response to Fe-limitation

To identify candidate taxa driving the differences in metabolic activity identified in the nanoSIP and exometabolomic results, we compared the microbiome composition between the two Fe conditions. Using 16S rRNA gene profiling at days 5 and 9, we found that the bacterial communities in the Fe-replete samples differed significantly from the Fe-deplete samples, with the greatest difference in composition at day 9 (Supplementary Fig. S6B). At day 9, we found that of the 39 bacterial ASVs detected in the community, four ASVs were significantly more abundant in the Fe-deplete conditions (Fig. 5A). This included an uncultured *Cryomorphaceae* ASV which dominated the Fe-deplete community, reaching up to ∼50% relative abundance (Fig. 5A). There was another set of ASVs that maintained similar relative abundance values regardless of Fe condition, which we designate as persistent taxa. This group included a *Roseibium* assigned ASV (10-15%, ASV5) and a *Rositalea* assigned ASV (∼3-5%, ASV8) which made up a significant portion of the iron-deplete community (Fig. 5A).

**Figure 5.**
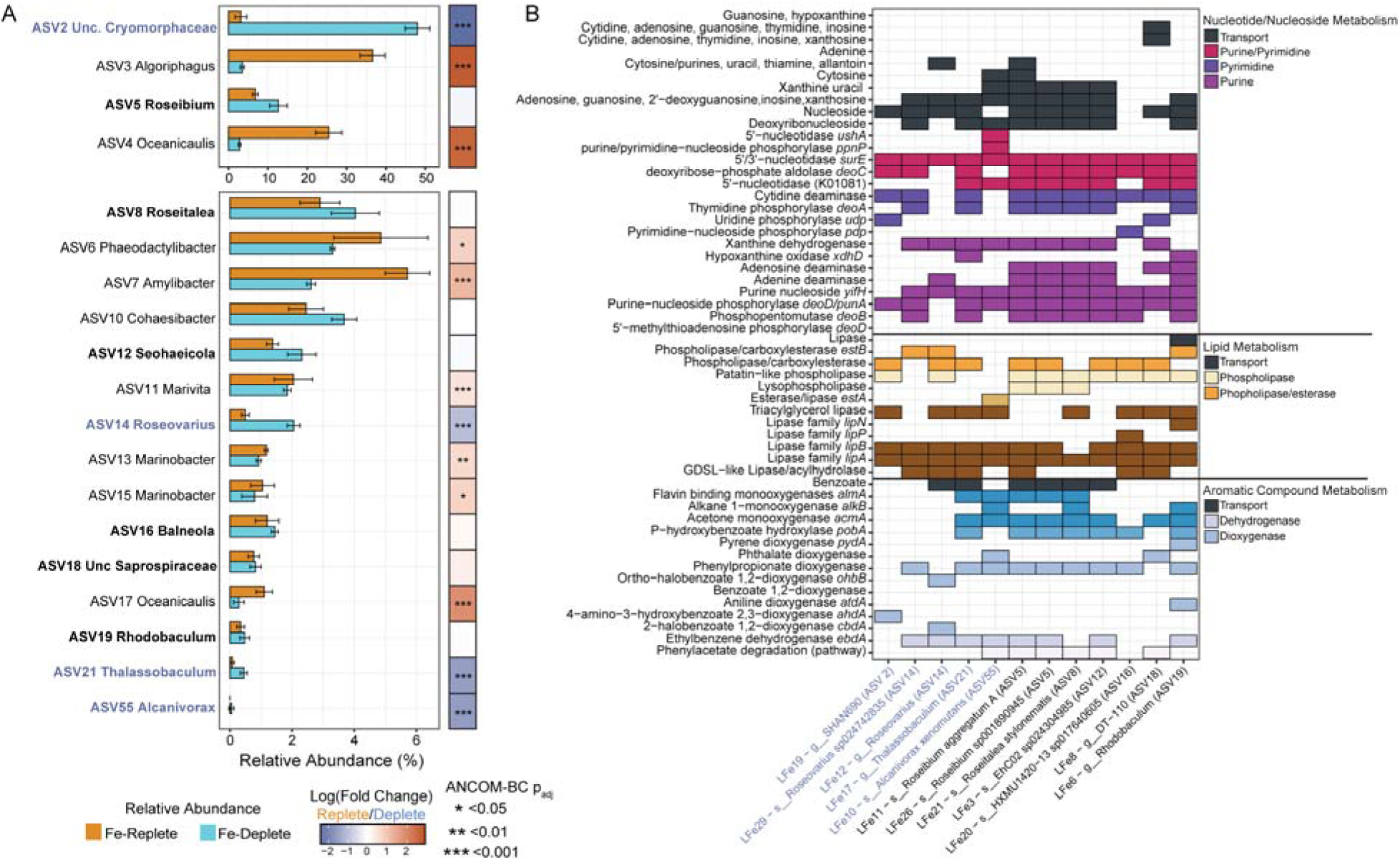
Taxon-specific response to Fe-limitation and their genomic potential to consume exometabolites enriched in the Fe-limited diatom exudate. **A)** 16S rRNA profiles of ASVs that are >=0.5% relative abundance in either Fe-replete or Fe-deplete day 9 cultures. The tileplot indicates results from the ANCOM-BC analysis. ASVs names that are bold indicate taxa significantly enriched (blue text) or persistent (no change, black text) in the Fe-deplete community. **B)** Presence/absence of key nucleobase metabolism, lipid metabolism, and aromatic compound metabolism genes or pathways in MAGs of taxa enriched (blue text) or persistent (black text) in the Fe-deplete conditions.

To determine whether these taxa had the capability to utilize the substrates identified from the exometabolomics analysis, we generated metagenome assembled genomes (MAGs) from the enrichment community. We found that the MAGs matching to the predominant ASVs in the Fe-deplete community contained genes indicating the ability to utilize lipids, aromatics and nucleosides and nucleotides (Fig. 5B). We recovered MAGs matching each of the four Fe-deplete enriched ASVs (2 matches for ASV14), and matches to six of the seven persistent ASVs (2 matches for ASV5, none for ASV10) (Supplementary Fig. S7). All 12 MAGs contained lipid catabolic genes, such as *lipA* (extracellular lipase gene) and *lipB* (lipase chaperone) used for lipid degradation (Fig. 5B). In addition, all MAGs contained genes involved in aromatic ring hydroxylating dioxygenases or monoxygenases, which are needed for aromatic compound breakdown (Fig. 5B). The *Thalassobaculum, Roseibium* and *Roseitalea* assigned MAGs were especially enriched in aromatic compound degradation, and they all carried transporters for benzoate (Fig. 5B). The MAGs were also enriched in genes to transport and catabolize purine and pyrimidine nucleotides/nucleosides and their precursors, such as genes encoding nucleotide phosphorylases, nucleotidases and deaminases (Fig. 5B). This indicates that the Fe-deplete MAGs have the genomic capability to utilize these compound classes, but they were not unique in this respect, because MAGs recovered from the taxa that were predominant in the Fe-replete condition (7 MAGs) also contained similar capabilities (Supplementary Table S10). This suggests that other factors aside from genomic content may contribute to their selective advantage under Fe-limitation, leading to their consumption of these compound classes.

Aside from these specific substrate utilization capabilities, the 12 MAGs also contained genes indicating the ability to use alternative sources of C in the form of inorganic C (CO_2_ /CO), and biopolymer C (CAZymes), and acquire Fe through multiple strategies (siderophore production and transport, ferric/ferrous transport, heme uptake and degradation) (Supplementary Fig. S8). The MAG matching the most abundant ASV in the Fe-deplete condition (LFe19-g_SHAN690, family *Cryomorphaceae*) encodes proteobacteriorhodopsin (Supplementary Fig. S8). Only three MAGs (two persistent taxa and one Fe-replete enriched) were predicted to biosynthesize siderophores, yet all MAGs had the capacity to transport siderophores (Supplementary Fig. S8). Sulfur metabolism genes were well-represented across these 12 MAGs, including genes involved in the transport and breakdown (*tauD*) of taurine, methionine transport, and cysteine/methionine metabolism.

To look more closely at bacterial Fe acquisition, we isolated and characterized one strain of the persistent taxa, a *Roseibium* sp. (strain C24; most closely related to *R. aggregata* via genome comparison, and ASV5 about 10-15% relative abundance under Fe-deplete conditions). The genome of *Roseibium* sp. C24 also possessed several siderophore biosynthetic pathways that could have allowed sequestration of Fe from the media (Supplementary Fig. S9). However, we found that while growth of *Roseibium* sp. C24 was not significantly impaired when grown in Fe-deficient minimal media, the Fe:Ca ratio was lower following 18h of growth (p<0.05; Supplementary Fig. S9). This indicates that this abundant taxon responded to Fe scarcity by lowering cellular Fe quotas, perhaps through metabolic compensation so growth rate was not impaired. The lower level of biomass Fe is consistent with the community-wide response determined by nanoSIMS single cell analysis (Fig. 1E), suggesting that some of the other persistent taxa might also be reducing their Fe quotas. Given this strain’s success under Fe-limitation (maintaining high relative abundance in the community), these results suggest metabolic flexibility with regards to Fe limitation could also provide a selective advantage in these communities, along with increased Fe scavenging.

## Discussion

In this study, we used nanoSIMS, nanoSIP, exometabolomics, and microbiome profiling to investigate how changes in diatom Fe nutritional status regulate the metabolic activity and microbial composition of the diatom phycosphere. We found that Fe-limitation led to the emergence of a population of bacteria with distinct metabolic activity and subsequent differences in carbon flow. The Fe-deplete cultures exhibited a shift to microbial consumption of metabolite classes including aromatics, purine and pyrimidine derivatives, and distinct lipid-like compounds. These results were complimented with the detection of Fe limitation-adapted taxa that can lower their Fe quotas and possess the genomic functional capabilities to utilize these metabolite classes.

Fe limitation shifted diatom exudation of aromatic exometabolites, which may partly explain the shift in carbon flow relative to the Fe-replete condition. Reactive oxygen species (ROS) accumulation often occurs under Fe-limitation in photosynthetic organisms due to impaired photosynthetic electron transport ^57,58^, leading to oxidative stress responses. Increased oxidative stress has been shown to increase antioxidant phenol exudation by *P. tricornutum* ^59,60^. It has also been shown that tryptophan synthase expression increased in Fe-limited *P. tricornutum*, which was hypothesized to be involved in the production of indole antioxidants to deal with oxidative stress while typical ROS defense proteins were down-regulated ^16^. Interestingly, tryptophan and aromatic amino acid synthesis markers have been shown to be upregulated in other Fe-limited diatoms ^61,62^, which may implicate tryptophan and other indole antioxidants as hallmark features in the exudate pool of diatom-dominated Fe-limited oceanic regions. The production of polyaromatic flavins likely also increases for Fe-starved diatoms due to the substitution of ferredoxin with flavodoxin and the increase in ferric reductase flavoproteins ^16^, consistent with our finding of increased lumichrome exudation.

Consumed metabolites associated with Fe-deficient conditions also included small aromatic compounds such as syringaldehyde, benzylideneacetone, lumichrome, tryptophan, and xylidine which likely favor the growth of aromatic specialists in the phycosphere. Xylene, an aromatic volatile organic compound (VOC), was shown to be exuded by *P. tricornutum* under light driven growth, possibly as a response to light stress ^63^. *Roseibium* was one of the specialist taxa that has been shown to consume aromatic and aliphatic hydrocarbon VOCs produced by *P. tricornutum* ^64^, and in this work was found to persist in high relative abundance in the Fe-limited microbiome. This taxon, along several other taxa enriched in the Fe-limited microbiome, possesses benzoate transport ability and, as previously mentioned, several genes needed to oxidize hydrocarbons. In earlier work we did not find that our isolate bacteria could use lumichrome as the sole carbon source for growth, but it did alter the *P. tricornutum* enrichment community composition when added exogenously ^27^. Lumichrome can act as a quorum-sensing mimic ^65^, which in this case could contribute to the stimulation of some of the bacterial taxa, such as *Thalasobaculum*, which increased in abundance under Fe-limitation and with exogenous lumichrome additions ^55^. Our study suggests more broadly that environmental factors which increase the exudation of aromatic compounds beyond VOCs may play a role in shaping the phycosphere community. Importantly, we demonstrate that these aromatic compounds can be putative resources for the microbiome when other sources have been processed. Many of Fe-limited taxa encode genes for use of diverse aromatic compounds which indicates that those with diverse metabolic flexibility may gain a competitive advantage during shifts in phototroph host physiology.

Fe limitation also resulted in an increase in the abundance and consumption of numerous purine and pyrimidine derivatives, indicating this was another important group of metabolites that could be implicated in changing carbon flow. Accumulation of hypoxanthine, thymine, uracil, and guanine may stem from increased nucleobase turnover caused by oxidative stress, or from reduced activity in iron-dependent recycling enzymes such as xanthine dehydrogenase^66,67^. Conversely, adenine was more abundant in Fe-replete axenic cultures, likely reflecting its additional role in the biosynthesis of central metabolites ATP, NADH, and SAM. Nearly all of the taxa enriched under Fe-limitation possessed the genomic capability to transport, degrade, and transform a variety of nucleotides, via nucleotidases, deaminases, and phosphorylases, supporting the importance of purine and pyrimidine-like compounds in the exometabolite pool. These metabolites reflect a general trend of greater exudation of aromatic organic molecules under Fe-limited conditions that are likely to be difficult and slow to break down by microbes relative to other major classes of exuded molecules such as peptides.

Lipid-like exometabolites were the most dominant stoichiometric group in both the diatom exudate and consumed fraction within the Fe-limited microbial community. It is likely that Fe deficiency led to increased lipid-like molecules in the exudate, as lipid accumulation has been documented in *P. tricornutum* undergoing nitrogen stress ^68^ and Fe-stress in *Chlamydomonas reinhardtii* ^69^. The average retention time was significantly greater across the Fe-limited exometabolites, consistent with an increase in the exudation and consumption of hydrophobic lipid-like molecules, including hydroxydodecanoate and aleuretic acid. These lipid-like compounds may serve as a valuable source of C for specialists such as *Alcanivorax*, which are known to specialize in hydrocarbon substrates, and are enriched in oil spills within the marine environment ^70,71^. We found that this taxon and several others were significantly enriched in the Fe-limited microbiome, and genomic analysis demonstrates lipid (*e.g.,* TAG lipase) and saturated hydrocarbon (*e.g.*, alkane monooxygenase) degrading pathways. Since lipids and aromatic DOM generally are slower to degrade and require specialist abilities, leading to slower turnover rates, it is possible that the usage of these resources reflect the low incorporation of recently fixed C in the active population of the Fe-limited microbiome.

Overall, we establish a systems-level understanding of how Fe-limitation impacts the model diatom, *P. tricornutum,,* the diatom-derived DOM pool,and the responding microbiome. By leveraging single-cell C incorporation measurements and molecular tools (*e.g.,* exometabolomics, metagenomics), we provide insight into how C and Fe availability control microbiome diversity and function. Indeed, it is known that C and Fe can co-limit bacterial growth and contribute to microbiome structuring in the ocean ^21,22,72,73^. This study more tightly constrains bacterial responses to changing C and Fe regimes, and, despite an overall reduction in DOC exuded by the diatom, we demonstrate that specialized taxa can persist and even increase in activity in response to diatom Fe limitation. Based on our collective results, we propose that DOM composition played a primary role in the microbiome response to reduced Fe. This metabolic response could be coupled to Fe-limitation, since instead of increased Fe scavenging, we observed an overall reduced Fe-quota in bacterial single cells, suggesting a switch to carbon metabolism requiring less Fe. Although this study focused on putative substrate exometabolites, there were many exometabolites that were produced in the presence of the microbiome, such as putative antimicrobials that could regulate species-specific microbial members under different Fe-regimes. Given similar genomic capacities for substrate usage in the MAGs from the different Fe treatments, we expect that future work examining transcriptional changes in the microbiome with Fe will further elucidate how Fe and C both work to shape microbiome responses to diatom Fe-limitation. Additionally, as exometabolite composition can vary based on host species^55^, more work to characterize exometabolomic responses with other phototrophs that persist in Fe-limited environments should be performed. Understanding how shifts in host exudate composition due to environmental condition impact the structure and biogeochemical feedback of heterotrophic bacteria will better constrain the fate of phototroph-derived carbon.

## Acknowledgements

This research was funded by U.S. Department of Energy (DOE) Office of Biological and Environmental Research grant SCW1039 under the microBiospheres Science Focus Area. The work at LLNL was performed under DOE auspices by Lawrence Livermore National Laboratory under Contract DE-AC52-07NA27344. We thank Kristina Rolison, John Casey, Ali Navid, and Ty Samo for their valuable feedback and discussions. We also thank Ty Samo for assistance in the calculations for total carbon content within single bacterial cells. LLNL-JRNL-2016953-DRAFT.

## Author Contributions

Conceptualization: N.E.G., R.B. and R.K.S. Methodology: N.E.G., X.M., and R.K.S. designed experiments. Investigation: N.E.G. performed Fe-limitation growth experiments. X.M, P.K.W., and C.R. performed nanoSIMS analyses, N.R.C and R.B. performed metabolomics, C.A.C and E.H. performed DOC analyses, N.M and P.D. performed bacterial isolate Fe-limitation analyses. Data curation/Formal analysis/Validation: N.E.G., J.A.K., R.B., P.K.W., X.M., P.D., E.H. Writing—original draft: N.E.G., R.K.S., P.K.W., X.M. R.B. Writing—review and editing: all authors

