## Supplementary Methods S1 for "Iron limitation alters diatom carbon flow through shifts in microbiome exometabolite consumption"

**Affiliations**

**File Includes:**

Supplementary Figures 1-9

Supplementary Methods

References

**Supplementary Figures**

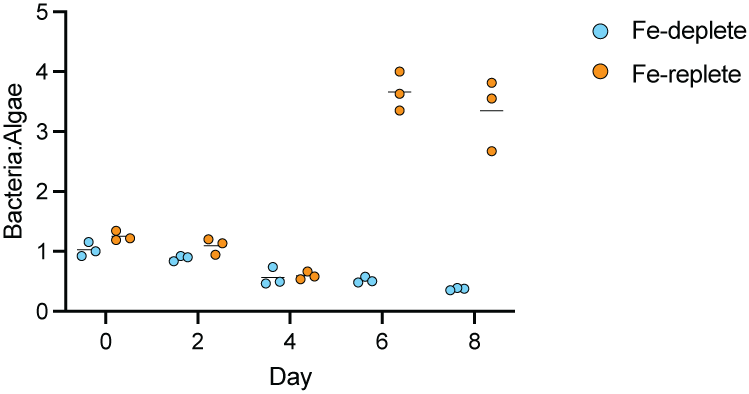

**Supplementary Figure S1. Ratios of bacterial cells to algal cells across time in the enrichment community.** Ratio calculated from algal and bacterial flow cytometric counts presented in Figure 1A–B. Treatments include Fe-replete (2 µM-added Fe) in orange and Fe-deplete (10 nM-added Fe) in blue. Points represent ratio from each replicate, and black line indicates mean between the triplicates.

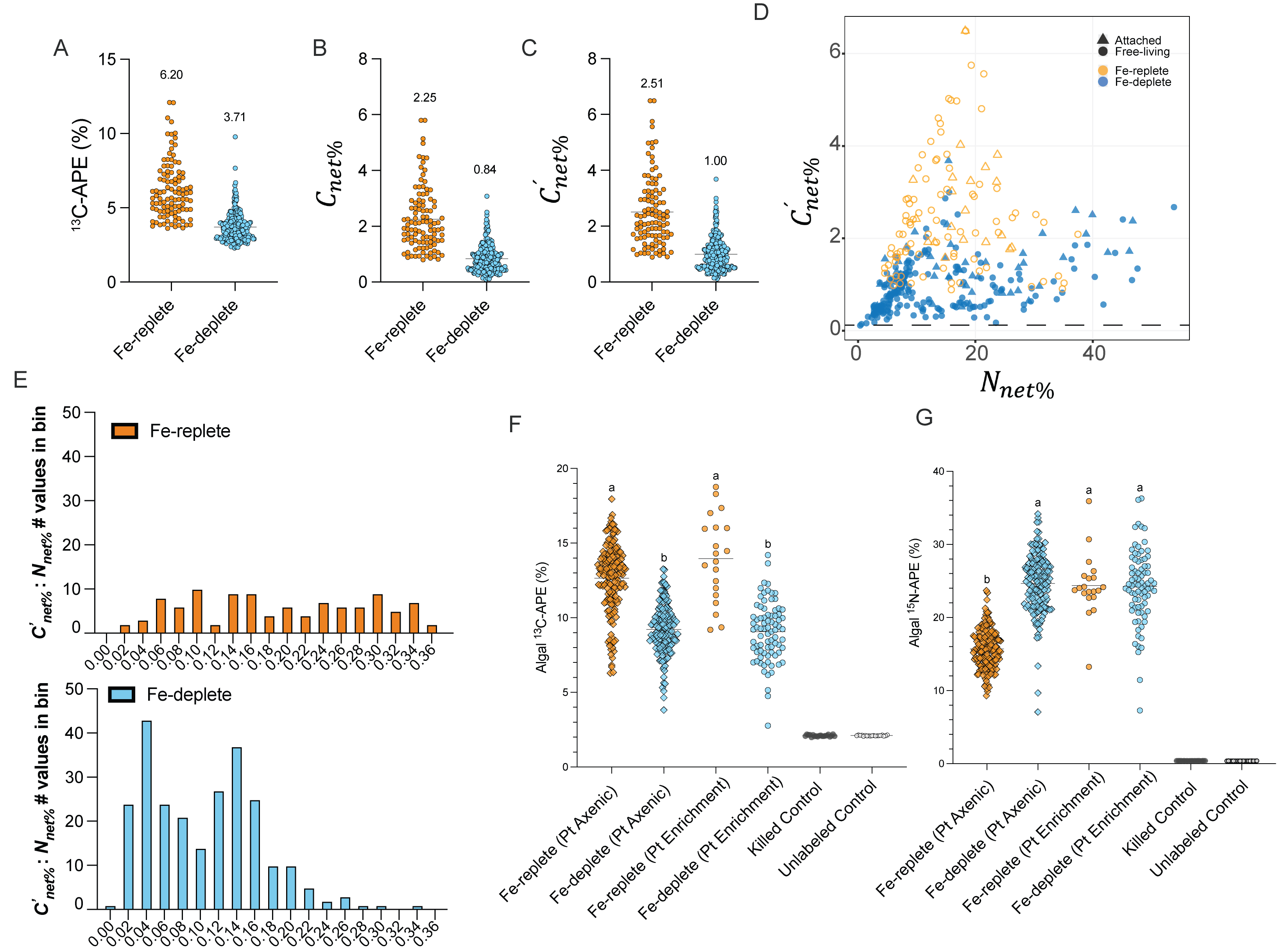

Supplementary Figure S2. **NanoSIP results**. Comparison of bacterial ^13^C incorporation values using the following calculations: A) APE [%]= (13C/12C+13C)*100 , B) $C_{net\%}$ using initial ^13^C input as 99%, C) $C_{net\%}^{´}$using initial ^13^C input as diatom ^13^C enrichment. Mean values are plotted above each treatment. (D) Bacterial $C_{net\%}^{´}$ vs $N_{net\%}$showing attached vs unattached bacteria. (E) Comparison of $C_{net\%}^{´}$ : $N_{net\%}$ distributions of Fe-replete (orange) and Fe-deplete (blue) values across all bacterial cells. Diatom atom percent excess (APE) of (F)^13^C-bicarbonate and (G) ^15^N-ammonium in *P. tricornutum* cells incubated under Fe-replete versus Fe-deplete conditions during the nanoSIP experiment. Triangles = axenic P. tricornutum cells, circles = P. tricornutum cells from the enrichment culture. Controls (killed and unlabeled) are shown indicating background values of ^13^C and ^15^N. Letters summarize the results of statistical comparison between Fe-treatment and culture was done using the Kruskal-wallace test with Dunn's multiple comparison. Groups are considered significantly different if p<=0.05.

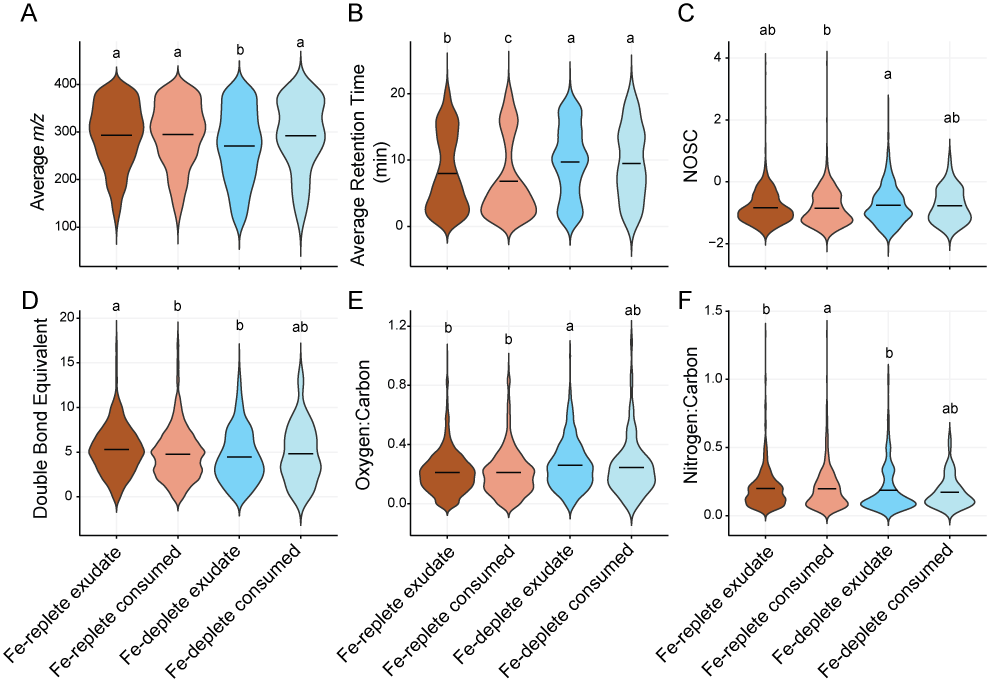

**Supplementary Figure S3.** Chemical characteristics of exometabolites derived from the CoreMS annotations. The distribution in chemical characteristics A) average mz B) average retention time C) nominal oxidation state of carbon [NOSC] D)double bond equivalent [DBE] E) oxygen to carbon ratio and F) nitrogen to carbon ratio across CoreMS annotated metabolites was assessed, comparing exometabolites significantly (p<0.05) enriched in the Fe-replete (n = 3086) or Fe-deplete (n= 1307) axenic P. tricornutum exudate and exometabolites significantly consumed in the Fe-replete (n= 2262) or Fe-deplete (n=193) enrichment cultures. Letters above each column represent results from a Kruskal Wallace + Dunns statistical test (p<0.05 is sig).

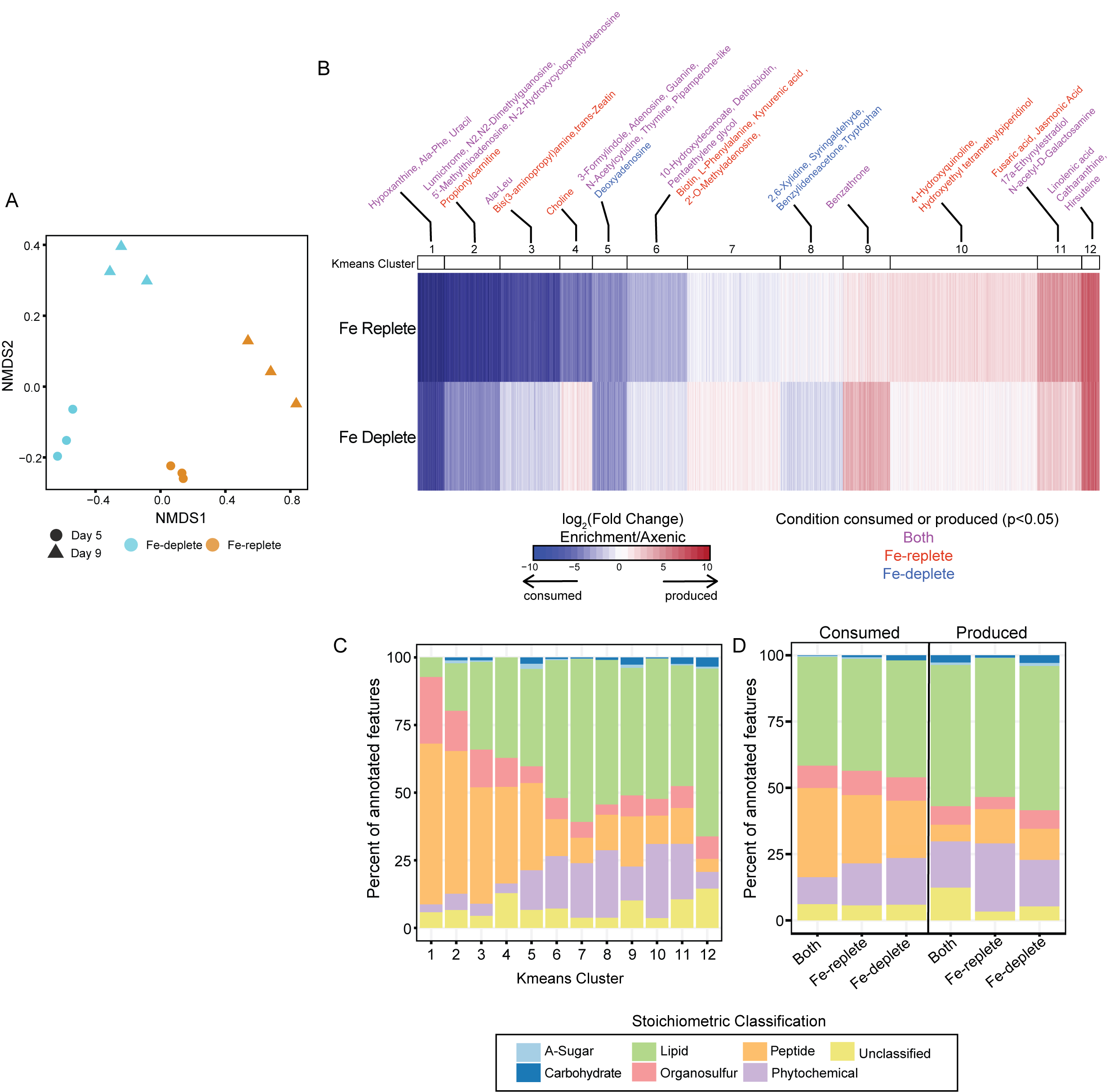

**Supplementary Figure S4. Patterns across all exometabolites.** A) Consumed or produced exometabolites differ based on iron condition. B) K-means clustered exometabolites based on log2(fold change) between the enrichment culture and axenic culture in the iron replete or iron deplete condition. Exometabolites are annotated in each cluster if the feature had a detected fragmentation spectrum and confidently matched to the reference metabolite. C) All features annotated using the CoreMS pipeline in each kmeans cluster, grouped by stoichiometric classification. D) Features that were significantly (p<0.05) consumed or produced, grouped by iron condition and stoichiometric classification.

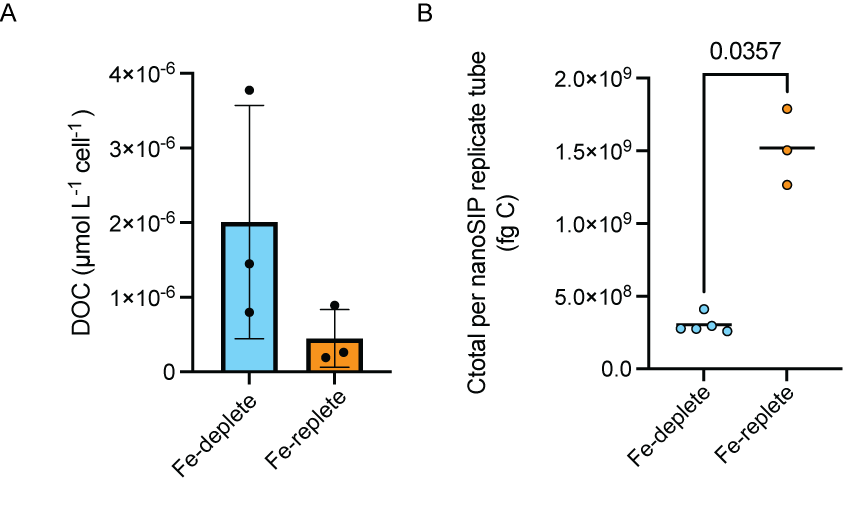

**Supplementary Figure S5. Additional carbon measurements.** A) Dissolved organic carbon (DOC) measurements, normalized to algal cell abundance, from enrichment cultures on Day 5. Bars represent mean and error bars represent one standard deviation between biological triplicates. There was no significant difference between Fe-condition. B) Calculated total carbon (Ctotal) incorporated into bacterial cells calculated based on net bacterial biomass production for individual growth experiments (number of bacteria x mass per cell x $C_{net\%}$) ^1^.

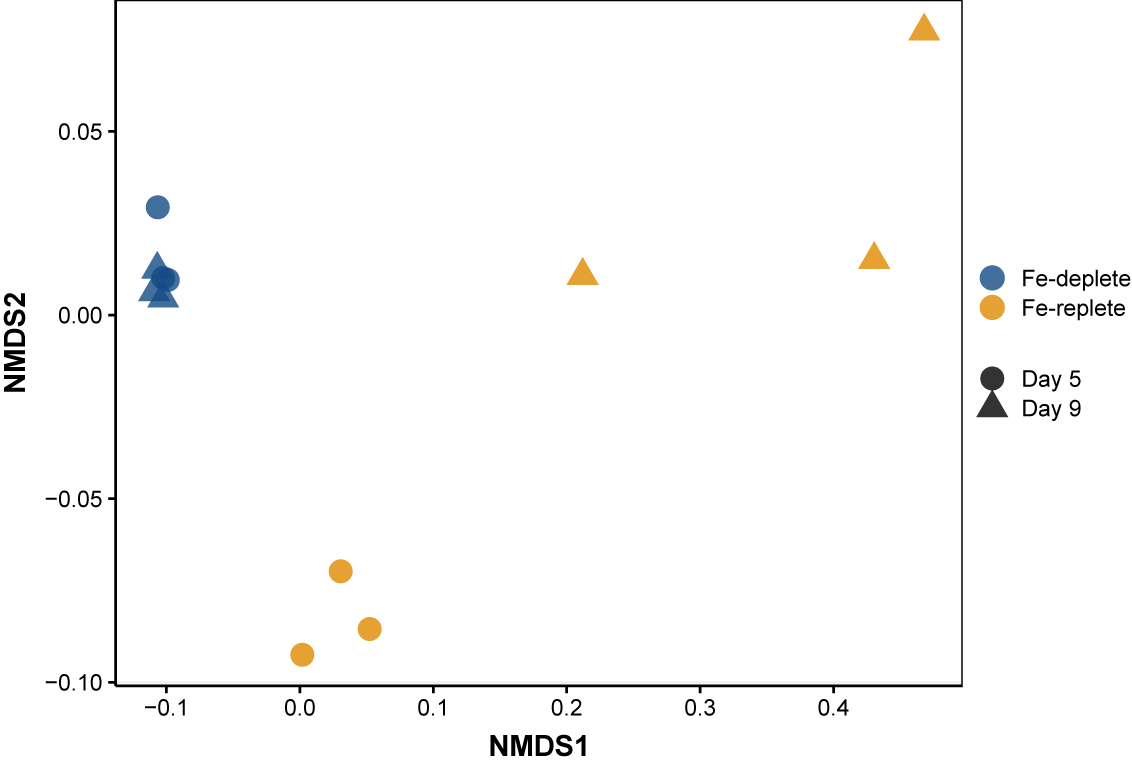

**Supplementary Figure S6.** Non-metric Multidimensional Scaling (NMDS) ordination of beta-diversity analyses at the ASV level in the 16S rRNA amplicon data.

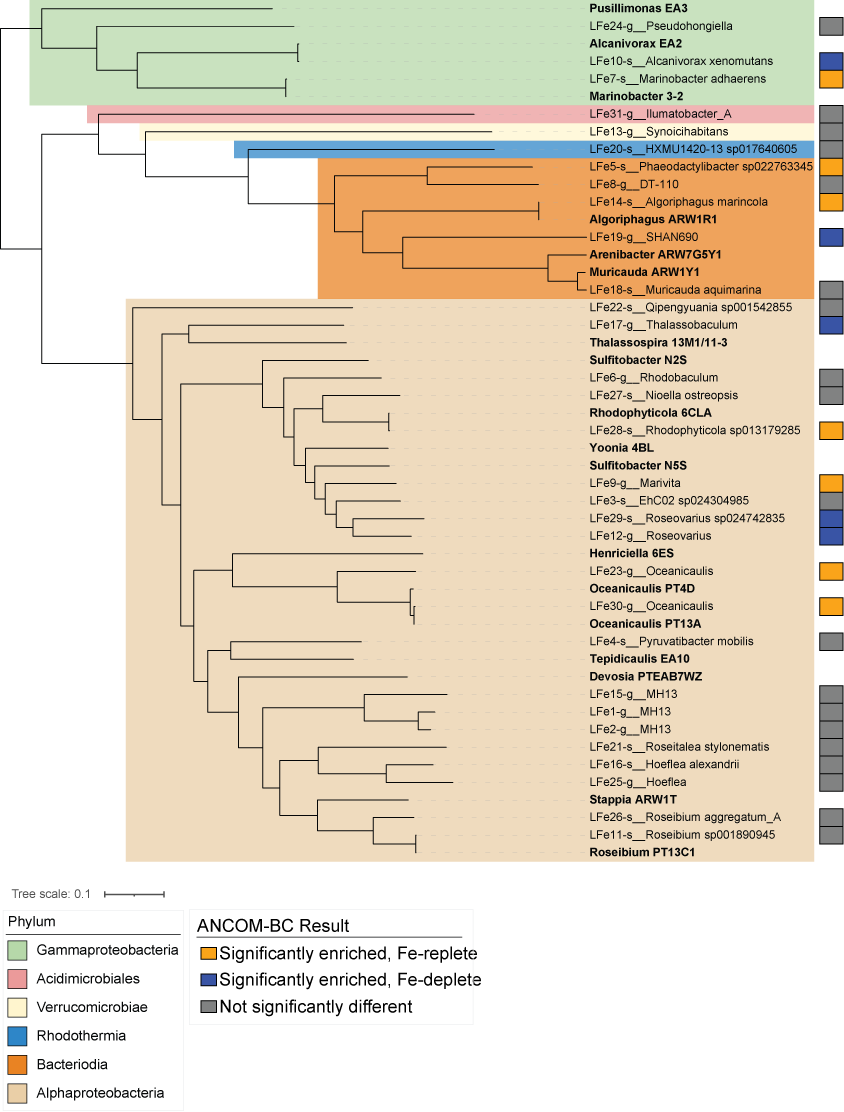

**Supplementary Figure S7. Phylogenetic relationship of MAGs recovered from *P. tricornutum* enrichment culture based on orthogroup analysis, color coded by phylum**. Bold text shows reference genomes from sequenced isolates obtained as previously described^2, 3^. Colored boxes on the right indicate whether the corresponding ASV in the 16S rRNA analysis was significantly differentially abundant in either Fe condition.

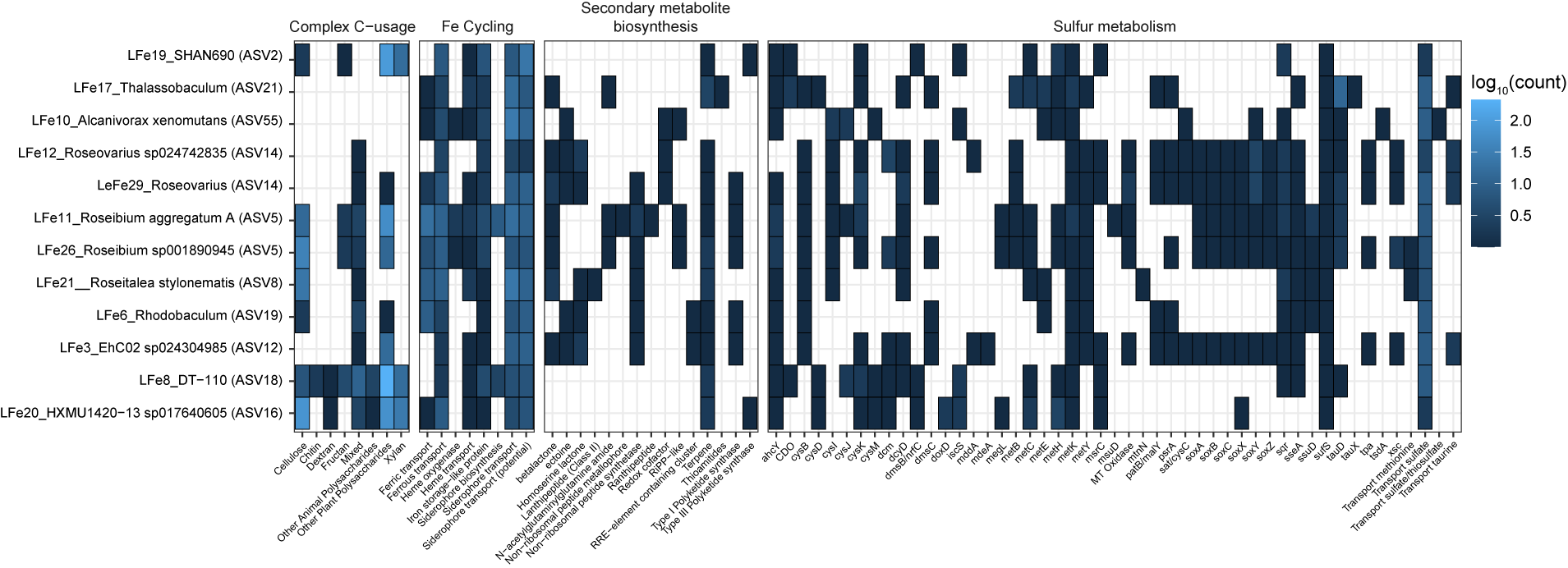

**Supplementary Figure S8.** Presence/absence of other pathways of interest across the Fe-deplete MAGS.

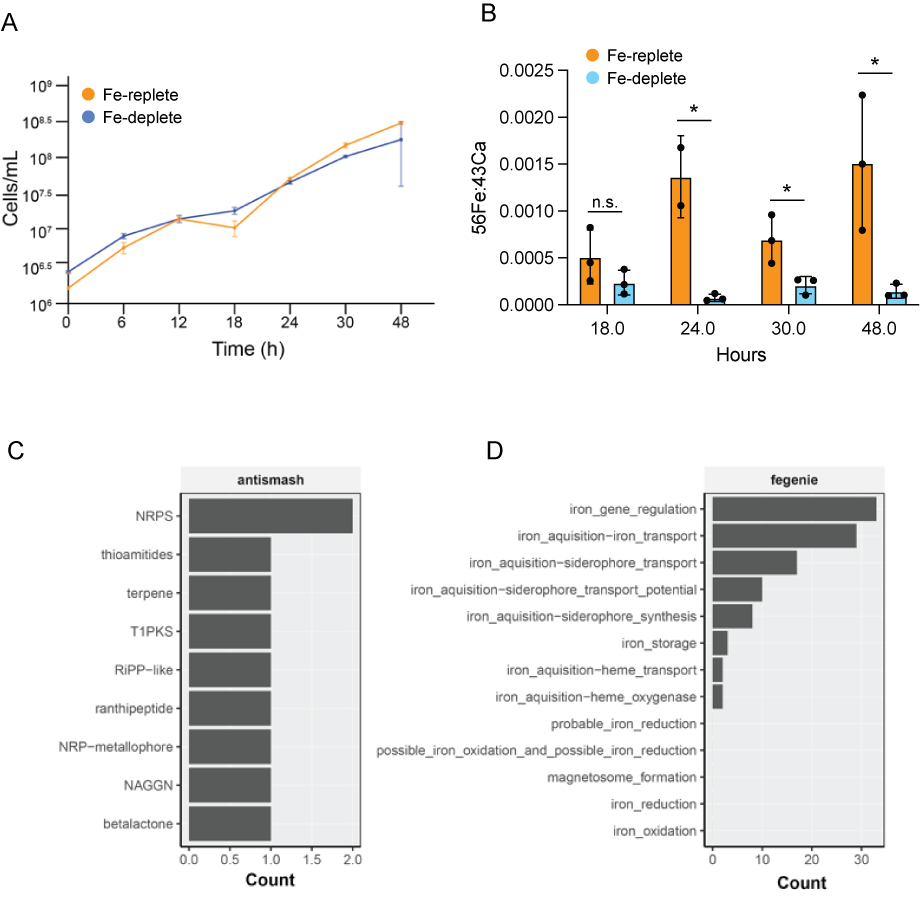

**Supplementary Figure S9**. **Growth response, Fe quota and secondary metabolite and Fe cycling capability of low-Fe isolate *Roseibium* sp. C24**. A) Growth over time of C24 in minimal media with (Fe-replete) or without added Fe (Fe-deplete). The points show the means and error bars show the standard deviation from the mean. B) ICP-MS derived mean Fe quotas of C24 grown under the Fe-deplete or Fe-replete condition at different time points of growth. Fe was normalized to Calcium after removal of background Fe values at T=0h. p-values calculated between Fe-replete vs Fe-deplete values using the students t-test at each time point. C) Genomic ability to synthesize secondary metabolites (antiSMASH) derived from genomic analysis of *Roseibium* C24. D) Fe transport and siderophore synthesis (FeGenie) capability of *Roseibium* C24.

**Supplementary Methods**

***Exometabolomics sample preparation***

Samples were solid phase extracted following standard methods ^4-6^. Briefly, 0.1 g PPL cartridges (Agilent Technologies) were primed using 3 mL LC-MS grade methanol (VWR Chemical), 3 mL 0.1% trace metal grade HCl (Fisher Chemical), and 3 mL ultrapure water. 42.5 mL of filtered spent medium were loaded onto the primed cartridges at their natural pH. Cartridges were then rinsed with 3 mL ultrapure water to remove any residual salts prior to elution in 1.5 mL LC-MS grade methanol. The resulting extracts were dried down to approximately 100 μL and rehydrated with ultrapure water to a final volume of 1 mL. Prior to analysis, 200 μL of the rehydrated extracts were transferred to 300 μL polypropylene HPLC vials (VWR) and spiked with 1 μM of an internal standard (cyanocobalamin, Sigma Aldrich) for quality control. To ensure reproducibility over the course of analysis, a pooled sample was generated by combining equal volumes of each sample and was analyzed several times during LC-MS analysis.

***LC-MS analysis***

Samples were injected (20µL volume) onto a C18 column (Phenomenex Biozen XB-C18, 1.7 µm particle size, 2.1x100 mm) held at a temperature of 30°C. The solvent flow rate was 0.3 mL/min with a gradient of water (solvent A) and methanol (solvent B), both containing 0.1% formic acid. The gradient increased from 5% to 95% solvent B over 20 minutes and was held at 95% B for 5 minutes before re-equilibration to initial conditions. The column oven was held at a temperature of 30°C. The mass spectrometer was equipped with a heated electrospray ionization (ESI) source in positive mode. ESI source parameters were set to a capillary voltage of 3500 V, sheath, auxiliary and sweep gas flow rates of 50, 10, and 1 (arbitrary units), and ion transfer tube and vaporizer temperatures of 325 °C and 350 °C. MS1 scans were collected over a m/z range of 100–1000 in positive mode. MS/MS fragmentation spectra were collected in data-dependent acquisition mode with a precursor quadrupole isolation width of 1.6, an acquisition rate of 35 spectra per second, an HCD energy of 30, and ion trap detection. Precursor ions were excluded dynamically for a duration of 5 s within a mass tolerance of 10 ppm.

***MS/MS matching in MS Dial***

MS/MS matching was based on an accurate mass tolerance of 0.005 Da for MS1 and 0.1Da for MS2 with default annotation cutoff parameters (dot product score of 600 and minimum of 3 matched peaks). For alignment of features across samples, an MS1 tolerance of 0.01Da and retention time tolerance of 0.5 min was used.

***Exometabolite molecular formula assignment***

For each data file, spectra were averaged over 1 minute time intervals from 0-25 minutes and mass calibrated based on a binomial correction function derived from a set of background polysiloxane masses. Formula were assigned with elemental stoichiometry of C1-50 H4-100 O0-20 N0-8 S0-2 Na0-1, within ranges of 0.3-3 for H/C, 0-1.2 for O/C, and 0-20 for double bond equivalents, ±2 ppm for mass error, and a charge state of +1.

***16S rRNA amplicon processing, sequencing and analysis***

Filters for DNA extraction were thawed on ice and placed into Lysing Matrix E, 2mL tubes (MP Biomedicals), 700 µL of Solution A (0.5% SDS, 20 mM sodium acetate, 10 mM EDTA, molecular grade water) was added, and the tubes were bead beat using the MP Biomedicals FastPrep-24 Bead beating system (40 s, level 5.5). Then, 500 µL of basic phenol:chloroform:isoamyl alcohol (pH 8.0) was added and the tubes were bead beat using the MP Biomedicals FastPrep-24 Bead beating system (40 s, level 5.5). The supernatant was transferred to a new tube and the aqueous layer re-extracted twice with 500 µL of pure chloroform. Then, sodium acetate was added to the final aqueous layer to 0.3 M concentration, 2-2.5x volumes of ice cold 100% ethanol added, and tubes were incubated at -80^o^C to precipitate the DNA for at least 1 h. The DNA was pelleted by spinning at maximum speed at 0^o^C for 1 h, then washed with 70 % ethanol and spun again for 15 min at maximum speed. The pellet was dried and resuspended in molecular grade water. The 16S rRNA gene was amplified from the extracted DNA using the Illumina V3-V4 universal primers 5'-TCGTCGGCAGCGTCAGATGTGTATAAGAGACAGCCTACGGGNGGCWGCAG-3’ (forward) and 5'-GTCTCGTGGGCTCGGAGATGTGTATAAGAGACAGGACTACHVGGGTATCTAATCC-3; (reverse), following the manufacturers protocol (IDT, Coralville, IA)^7^. The PCR products were confirmed by gel electrophoresis, diluted to 10 ng/µL, and purified and sequenced using the Illumina MiSeq 16SV4 procedure at Laragen Inc (Culver City, CA).

The raw data was processed using DADA2 v1.28.0^8^ in R (R Core Team 2013) v4.3.1 to assess read quality, trimmed and filter reads, remove chimeras (removeBimeraDenovo), and generate an amplicon sequence variant (ASV) table. Taxonomy was assigned using the RDP database (v18) and SILVA (v138). Phyloseq v1.44.0^9^ was used for statistical analysis of the 16S ASVs. Chloroplast and mitochondria sequences were removed, and ASVs that were not detected with at least 5 counts across at least 3 samples were filtered out to remove low prevalence ASVs^10^. ANCOM-BC v. 2.2.22^11^ with false discovery rate p-value correction (alpha = 0.05) was used for differential abundance testing of ASVs, comparing the Fe-replete to Fe-deplete conditions separately for each timepoint. The VEGAN package v.2.6-4^12^ was used to construct nonmetric multidimensional scaling (NMDS) plots using Bray-Curtis distances.

***Metagenome processing, sequencing and analysis***

To obtain metagenome assembled genomes (MAGS) of taxa enriched in the Fe-limited community, 50 mL of the low Fe acclimated enrichment culture was filtered onto a 0.2 µm Supor filter and total DNA extracted following the phenol:chloform protocol used for 16S amplicon analysis. Metagenome sequencing was done through the Joint Genome Institute (JGI) using the Illumina NovaSeq S4. Initial quality control was done by JGI using the JGI Standard Operating Procedure^13^. *P. tricornutum* reads were removed by mapping the quality-controlled reads against three *P. tricornutum* genomes downloaded August 2022 (NCBI Accessions ASM15095v2, HQ840789.1 and NC_008588.1) using Bbduk from BBMap v39.08^14^. The remaining bacterial reads were assembled using metaSPAdes v3.15.3^15^ in Kbase^16^. Contigs were binned using MaxBin v2.2.4^17^, MetaBAT v1.7^18^ and CONCOCT55 v1.1.0^19^. CheckM v1.1.3^20^ was used to determine MAG quality and only MAGs with at least 75% completion and <25% contamination were retained for dereplication. dRep v3.4.2 ^21^ was used to dereplicate the MAG bins and Prodigal v2.6.3^22^ was used for gene calling.

Genes were annotated using Eggnogg-mapper v 2.1.12 ^23^, TransAAP with TransportDB 2.0 ^24^, FeGenie v1.2^25^, antiSMASH V7 ^26^, dbCAN2^27^, and GATOR (https://github.com/jeffkimbrel/gator). Details on gene annotation are described in Supplementary Method S1. Orthofinder 2.5.5^28, 29^ was used for orthogroup identification between assembled MAGs from this study, the *Roseibium* strain isolated from the low Fe enrichment, and genomes of bacteria isolated from *P. tricornutum* previously^3^. The species tree from Orthofinder was visualized and annotated using iTol v5^30^. MAG taxonomy was assigned using the Kbase application “Classify Microbes with GTDB-tk-v2.3.2” r207 release using default parameters^16^.

***Flow cytometry***

Diatom and bacterial abundance were assessed using an Attune benchtop flow cytometer with a CytKick autosampler (Thermo Fisher Scientific, Waltham, MA) fitted with a blue laster (excitation 488 nm). One mL of sample was collected and immediately fixed with glutaraldehyde (0.25% final concentration), frozen and stored at −80 °C. Fixed samples were diluted in sterile-filtered media to achieve >4000 event counts per second and stained with 1X SYBR Gold (Thermo Fisher Scientific, Waltham, MA) for 10 minutes in the dark prior to analysis. The instrument parameters were set up to threshold at 0.1 x 1000 on the BL1 detector, sample acquisition volume of 200 µL run at a flow rate of 100 µL/min. Gating was performed based on forward scatter, side scatter, and fluorescence of SYBR Gold (BL1 detector, emission/BP of 530/30). MilliQ blanks were run between treatment replicate sets to reduce carryover, and media blanks were included to correct for background particle noise.

***Metal analysis by ICP-MS***

Metal concentrations were quantified using an Agilent 7850 ICP-MS instrument (Agilent Technologies) equipped with a microplate adaptor. Measurements were conducted in helium (He) reaction gas mode to minimize interferences from doubly charged ions, oxides, and polyatomic species. Instrument operating conditions included an RF power of (1550) W, a nebulizer gas flow rate of (1.08) L min⁻¹, and a dwell time of (0.1) ms per isotope. Calibration standards (0, 0.1, 1, 10, 100, and 1000 ppb) were gravimetrically prepared from commercial rare earth element TraceCERT® stock solutions and Inorganic Ventures ICP-MS grade standards. Milli-Q water blanks and acid matrix blanks were included as controls and analyzed every 20 samples to monitor background levels and instrumental drift. Samples were diluted to a final matrix of 1.5% (v/v) HNO₃, prepared using 0.2 μm-filtered Milli-Q water and ultrapure HNO₃ (OPTIMA™, ~70% w/w, ≥99.999% trace metal purity; Fisher Scientific, S020101TFIF01). A 1 ppb rhenium internal standard (Inorganic Ventures, CGRE1) in 1.5% HNO₃ was continuously introduced during analysis for signal normalization. Instrument tuning and performance optimization were conducted using a 1 ppb Agilent tuning solution (5185–5959). The sample introduction system was rinsed with 0.2 μm PES-filtered Milli-Q water between analyses. Quantification was performed using external calibration curves generated by linear regression of standard concentrations versus internal-standard-corrected signal intensities, with acceptable calibration defined as R² ≥ (0.999). Sample concentrations were calculated automatically using Agilent MassHunter software (version 5.2 Agilent Technologies). Data are reported as mean ± standard deviation of three independent sample preparations. No additional statistical tests were applied unless otherwise noted. To account for residual Fe possibly bound to the cell pellet in the Fe-replete condition, the averaged 56Fe value (0.967 ppm) from the start of the experiment (T0h) was subtracted from the final 56Fe values in the Fe-replete samples from the 18h, 24h, 30h, and 48h timepoints before calculating Fe:Ca (56Fe/43Ca).

***Metagenome assembled genome gene annotation***

Eggnogg-mapper v 2.1.12 ^23^ was used for initial functional annotation using DIAMOND (-m diamond) alignment ^31^ and an evalue threshold of 1e-10. Transporters were further annotated using TransAAP with TransportDB 2.0 ^24^. Genes involved in Fe cycling were identified using FeGenie v1.2 ^25^ and biosynthetic gene clusters were identified using antiSMASH V7 ^26^. Carbohydrate active enzymes (CAZYmes) Hidden Markov models (HMMs) were queried using dbcan12 with dbCAN2 ^27^. GATOR (https://github.com/jeffkimbrel/gator) was used to assess the metabolic pathway completeness for curated pathways relevant to phycosphere microbes.
